# Food restriction engages prefrontal corticostriatal cells and local microcircuitry to drive the decision to run vs conserve energy

**DOI:** 10.1101/2020.10.16.342808

**Authors:** Adrienne N. Santiago, Emily A. Makowicz, Muzi Du, Chiye Aoki

## Abstract

Food restriction (FR) evokes running, which may promote adaptive foraging in times of food scarcity, but can become lethal if energy expenditure exceeds caloric availability. Here, we demonstrate that chemogenetic activation of either the general medial prefrontal cortex (mPFC) pyramidal cell population, or the subpopulation projecting to dorsal striatum (DS) drives running specifically during hours preceding limited food availability, and not during *ad libitum* food availability. Conversely, suppression of mPFC pyramidal cells generally, or targeting mPFC-to-DS cells, reduced wheel running specifically during FR and not during *ad libitum* food access. Post-mortem c-Fos analysis and electron microscopy of mPFC layer 5 revealed distinguishing characteristics of mPFC-to-DS cells, when compared to neighboring non-DS projecting pyramidal cells: 1) greater recruitment of GABAergic activity and 2) less axo-somatic GABAergic innervation. Together, these attributes position the mPFC-to-DS subset of pyramidal cells to dominate mPFC excitatory outflow, particularly during FR, revealing a specific and causal role for mPFC-to-DS control of the decision to run during food scarcity. Individual differences in GABAergic activity correlate with running response to further support this interpretation. FR enhancement of PFC-to-DS activity may influence neural circuits both in studies using FR to motivate animal behavior and in human conditions hallmarked by FR.

## INTRODUCTION

Adaptive behavioral modification in times of food scarcity can provide individuals with an evolutionary advantage. In many diverse species, food restriction (FR) evokes an increase in running, which promotes adaptive foraging behaviors (de Lartigue G and M McDougle 2019; Södersten P et al. 2019). While running to seek new locations with potentially greater resources may be adaptive in some circumstances, FR-evoked running is potentially life threatening, if energy expenditure exceeds caloric availability. In adolescent mice, restriction of food to a two-hour window is non-lethal (Chowdhury TG et al. 2013; Chen YW, H Actor-Engel, et al. 2018), yet when given access to a running wheel, as in the activity-based anorexia (ABA) model (Gutierrez E 2013), FR mice can lose between 20-25% body weight in as little as three days, increasing risk for mortality (Chowdhury TG et al. 2013). The greatest increase in excessive running occurs during the hours leading up to the scheduled feeding time (food anticipatory activity, FAA)(Mistlberger RE 2020). The extent of FR-evoked running is hallmarked by variance, and individual differences in running responses correlate with an increase in anxiety-like behavior (Wable GS et al. 2015). Despite the prevalence of FR as a tool to motivate animal behavior (Goltstein PM et al. 2018), little is known about the neural pathways that drive FR-evoked hyperactivity.

In dorsal striatum (DS), where dopamine signals a feeding-entrained oscillator (de Lartigue G and M McDougle 2019), FAA requires expression of dopamine D1 receptors (D1R) (Gallardo CM et al. 2014). Striatal D1R cells are the major players in the direct pathway to initiate movement (Gerfen CR and DJ Surmeier 2011). FR increases DS sensitivity to D1R-mediated dopaminergic activity (Carr KD 2002). The pleasurable act of voluntary wheel running (Meijer JH and Y Robbers 2014) also induces a hyper-dopaminergic state in DS, wherein even aversive stimuli, such as an energy-depleted state, can activate circuitry of reward and addiction (Kanarek RB et al. 2009; Greenwood BN 2019). This positions DS as a likely target for executing FR-evoked running. The question remains: Is top-down control of DS-mediated running FR-dependent?

One major excitatory input to the DS, the medial prefrontal cortex (mPFC), has never been causally implicated in generating FR-evoked running. FR shifts circadian c-Fos expression in the mPFC to produce a spike in activity corresponding to FAA (Angeles-Castellanos M et al. 2007). mPFC c-Fos levels also increase during states of food reward expectation (Valdes JL et al. 2006). Temporal control of goal-directed behavior requires mPFC pyramidal cells expressing D1R (Narayanan NS et al. 2012), a cell type known to project from layer 5 of mPFC to DS (Anastasiades PG et al. 2019). We therefore used multiplexed chemogenetics to test the prediction that FR gates mPFC pyramidal cell control of hyperactive running. Specifically, we determined whether activation and suppression of these cells modulates FR-evoked running without altering running under *ad libitum* food. We further tested whether modulation of mPFC-to-DS projecting pyramidal cells is sufficient or requires additional mPFC pyramidal cells with diverse projections to modulate FAA running.

How does microcircuitry of the mPFC contribute to FR-evoked running? mPFC parvalbumin-positive GABAergic interneurons (GABA-INs) increase firing during goal-directed behavior (Kim H et al. 2016) and express D1R (Anastasiades PG *et al.* 2019). Likewise, the lateral hypothalamus, which registers and engages responses to FR and other aspects of homeostasis (Stuber GD and RA Wise 2016), projects directly to mPFC GABA-INs (Stuber GD et al. 2011). Moreover, lengths of GABAergic axon terminals targeting mPFC pyramidal cells correlate negatively with FR-evoked running (Chen YW et al. 2016), suggesting that mPFC GABA-INs may dampen FR-evoked running. Here, we ask whether the extent of mPFC pyramidal cell recruitment of GABA-INs contributes to individual differences in FR-evoked running. To this end, we quantified the extent of c-Fos expression in GABA-INs following excitation of mPFC pyramidal cells. We then used electron microscopy to ask whether GABA-INs target the mPFC-DS subpopulation and non-DS projecting pyramidal cells equally, or skews to favor one subpopulation. Our findings led us to a new interpretation of the role of GABA-INs in facilitating mPFC-to-DS pyramidal cells to drive running specifically during FR.

## METHODS

### Experimental design

Because of the prevalence of Anorexia Nervosa (AN) in adolescent females (Murray SB et al. 2017), and the relevance of this work to researchers studying that disorder, only adolescent female C57/BL6 mice were used for this study. Adolescent rodents are also more susceptible to FR-evoked running, vs adults (Gelegen C et al. 2006; Gilman TL et al. 2019), which parallels the common adolescent onset of AN in the human population (Herpertz-Dahlmann B 2015).

For all studies, mice were assigned randomly to receive injection of either control virus or viruses for transcription of Designer Receptors Exclusively Activated by Designer Drugs (DREADDs) before any measure of activity was collected. Power analysis was conducted to estimate the sample size needed.

To drive both cell firing and cell suppression within the same target population, we simultaneously injected two DREADD viruses into a single mouse, with the hM3D(Gq) receptor to activate cell firing and the kappa opioid DREADD (KORD) receptor to suppress cell firing. Because the Gq receptor ligand CNO can reverse-metabolize to clozapine to produce off-target effects (Gomez JL et al. 2017), we chose to use the C21 ligand, which has been extensively characterized to show minimal off-target agonist activity at the dosage we used (1 mg/kg) (Thompson KJ et al. 2018). The KORD receptor ligand, Salvinorin B (SalB) has also been characterized extensively to be pharmacologically inert except as agonist to KORD (Vardy E et al. 2015; Marchant NJ et al. 2016). To target mPFC pyramidal cells, we injected DREADD viruses under a CaMKIIα promoter into the mPFC (CaMKII group; Fig. 1A). To target the mPFC-DS pathway, we injected cre-dependent DREADDs into the mPFC and a retrograde cre virus into the DS (mPFC-DS group; Fig. 1A). mPFC-DS and CaMKII group experiments were run concurrently.

**Figure 1.**
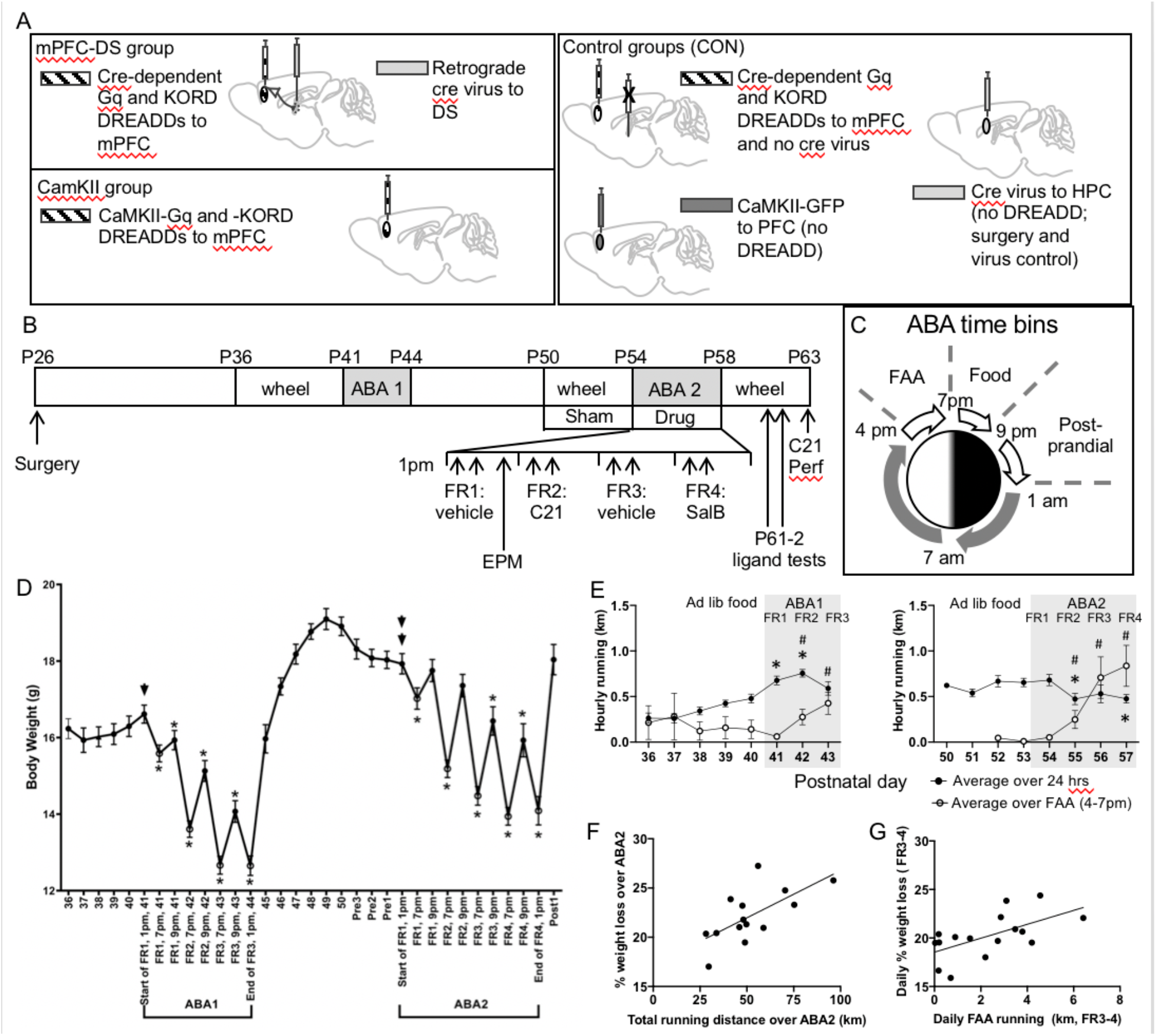
Experimental design and changes in body weight and wheel running evoked by ABA induction. **A**. Experimental groups consisted of 1) the mPFC-DS group, where cre-dependent Gq and KORD DREADDs were injected into the mPFC and retrograde cre virus was injected into the DS. This enabled selective DERADD expression in the subset of mPFC pyramidal cells that project to DS. 2) CaMKII group where viral transcription for Gq and KORD DREADDs was under control of a CaMKIIα promotor and delivered into the mPFC to target mPFC pyramidal cells. The control (CON) group consisted of three subgroups: 1) ‘no-CRE-CON’ (N=6) were injected with cre-dependent DREADD virus in the mPFC without injection of the retrogradely transported cre-virus, thus ensuring that the cre-dependence of the DREADDs was functional 2) ‘GFP-CON’ (N=5) were injected with CaMKIIα-GFP virus into the mPFC for GFP expression selectively in mPFC pyramidal cells 3) ‘No-Drug-cre-CON’ subjects (N=5), were injected with GFP-cre virus into a control region, the dorsal hippocampus, and unlike the other 11 control subjects, they did not receive DREADD ligand to test for off-target effects of the ligand **B**. ABA induction schedule. All animals underwent viral injection surgery at or around postnatal day 26 (P26). After surgery, mice were singly housed. On P36±1, a wheel was introduced to the cage and baseline wheel running and weight measures were recorded. The first bout of ABA (ABA1) began at 1 pm on P41, at which time food was removed, but the wheel was left in cage. For a total of 3 days, food access was restricted to the hours of 7 pm to 9 pm, corresponding to the beginning of the dark phase of the light-dark cycle. At 1 pm on P44±1, food restriction of ABA1 was terminated: wheels were removed and food access was *ad libitum*. Following ~6 days of recovery, wheel re-acclimation began, followed by 4 days of baseline wheel running measurements. The second bout of ABA (ABA2) began at 1 pm, whereby food was removed at all hours except for 7 pm to 9 pm. Of the 4 food restricted days (FR1 through 4), the ligand, C21 for Gq-DREADD was injected on FR2 and the ligand, Salvinorin B (SalB) for KORD was injected on FR4, with vehicle injected on FR1 and FR3. Following 4 days of food restriction, food access returned to be *ad libitum* and wheel was available, so as to measure DREADD ligand modulation of wheel activity in the absence of FR. Following these ligand tests during recovery, animals were injected with either C21 or SalB, then euthanized approximately 2 hours later by transcardial perfusion under urethane anesthesia for retrospective analysis of neuronal activation based on c-Fos-immunoreactivity and for electron microscopic analyses. **C**. Schematic of the three time bins measured daily during FR and recovery: FAA (food anticipatory activity period, 4pm-7pm), food allowance period (Food, 7pm-9pm), and the post-prandial period (9pm-1am). **D.** Body weights across the days preceding, during and following ABA inductions of control (CON) mice. Single arrow points to the baseline value at the start of FR1 of ABA1, at 1 pm on P41. Double arrow points to the baseline body weight at the start of FR1 of ABA2. Asterisks indicate body weights that were significantly less than the baseline value. Values are mean ± SEM. **E**. Hourly wheel running of CON mice during the days preceding and during ABA. Filled circle: hourly wheel running distance averaged over the course of the 24 hour day; asterisk indicates significant difference vs measure from day prior to FR induction. Open circle: hourly wheel running distance averaged over the 3 hour FAA time bin (4pm-7pm); hashtag indicates significant difference vs measure from FR1, which occurs 3 hours following FR induction. Note the precipitous rise of FAA during ABA1 and ABA2. **F.** Total running over the four days of ABA2 correlated positively with percent weight loss over the same timeframe (p = 0.010, R=0.66). **G.** FAA running, averaged over FR3 and FR4, correlated positively with daily percent weight loss averaged over the same period (p = 0.014, R = 0.602).

Littermates of mPFC-DS and CaMKII subjects were used as control subjects for both experimental groups. Three different types of control subjects were used, all lacking expression of DREADD genes (Fig 1A). ‘GFP-CON’ mice (N=5) were injected with CaMKIIα-GFP virus into the mPFC. This group controlled for energy expenditure required to replicate the virus in pyramidal cells of the mPFC. ‘No-cre-CON’ subjects (N=6) were injected with cre-dependent DREADD virus in the mPFC without injection of the retrogradely transported cre-virus. This control was used to ensure that the cre-dependence of the cre-dependent DREADDs was functional, and was validated based on the finding that neither DREADD reporter was transcribed for these controls. Both the GFP-CON and No-cre-CON groups, (N=11, collectively) received DREADD ligand administration that was identical to that of experimental DREADD mice, and were perfused alongside mice from the CaMKII and mPFC-DS experimental groups. A third control group, ‘No-Drug-cre-CON’ subjects (N=5), were injected with GFP-cre virus into a control region, the dorsal hippocampus, to control for surgical experience and energy expenditure required to replicate cre virus within brain. In contrast to the other control groups, the No-Drug-cre-CON subjects did not receive DREADD ligand, so as to provide a control to assess whether drug administration in the absence of DREADD receptor would perturb behavior. These mice were perfused immediately after the end of the second round of FR. We detected no significant differences between control groups (see Results section) for any of the variables we observed in this report, and so data for all three types of control were pooled under the ‘CON’ group (N=15). This CON group was used as comparison for all data points collected from the first introduction of the wheel to the end of the second bout of FR. For data points collected during recovery from the second bout of FR, only the mice from the GFP-CON and No-cre-CON groups were used, as the No-Drug-cre-CON subjects were perfused immediately following the second bout of FR.

Fifteen subjects were used for the CaMKII group. Of these, 8 received C21 on the second day of FR (FR2) and 10 received SalB on the fourth day of FR (FR4), with 3 mice receiving both C21 on FR2 and SalB on FR4. For the mPFC-DS group, all 9 mice received C21 to activate cell firing on FR2 and received SalB to suppress cell firing on FR 4. Our wheel hub malfunctioned on FR2 for one cohort consisting of two ‘no-cre CON’ (described below) and two mPFC-DS mice. This warranted exclusion of data from time-bins overlapping with the outage, but not exclusion of data at points when the hub was functioning. Otherwise, exclusion criteria were limited to death, abnormal locomotion on the elevated plus maze (administered before DREADD ligand drug administration), and insufficient bilateral viral replication in the region of interest (2 animals, with one each from the CaMKII and mPFC-DS groups).

Human bias was limited by masking group identity to keep the experimenter blind to experimental condition during all feeding, weight measurement, and drug/vehicle administration procedures. Wheel count was automatized, and analysis of immunofluorescent images, used to assess DREADD modulation of neurons, was also performed blind to experimental condition.

#### ABA induction

Because AN commonly has adolescent onset and a relapsing course (Wentz E et al. 2009), all mice (both control and experimental) underwent two bouts of ABA (FR plus wheel access), timed during mid and late adolescence (Fig. 1B), as previously described (Chen YW *et al.* 2016; Chen YW, AD Sherpa, et al. 2018). At P36±1, mice were provided with unlimited access to a running wheel capable of collecting continuous wheel count data (2654.86 turns per kilometer; Med Associates, Fairfax, VT). Following 5 days of acclimation, animals underwent a first bout of FR (ABA1), which lasted 3 days. During this time, food was restricted to a 2-hour period from 7pm to 9pm each night, unlimited in amount, while wheel access remained unlimited (Fig. 1C). After the third day of FR, the wheels were removed for a four-day recovery period. On P50±1, the wheels were replaced for 4 days of re-acclimation, after which the mice were subjected to a second bout of ABA lasting 4 days (ABA2). Afterwards, mice remained with unlimited wheel and food access until euthanasia by transcardial perfusion, 5 days later. One exception was of the 5 ‘No-Drug-cre-CON’ mice that were perfused at the end of ABA2, rather than euthanizing them 5 days after the end of ABA2.

#### Drug delivery

Subcutaneous injection of C21 was delivered on FR2, SalB was delivered on FR4, and vehicle was delivered on FR1 and FR3. On each of the four days leading up to FR1 mice were given a sham injection (no needle insertion), to climatize mice to the stress of brief restraint. Because C21 drug activity (Chen X et al. 2015; Thompson KJ *et al.* 2018) lasts longer than that of SalB (Vardy E *et al.* 2015), drug delivery times were staggered, aiming to target the hours of FAA and food availability. For the CaMKII experiment, C21 was delivered at 12 noon and 6:00pm, while SalB was delivered at 3:30 and 6:30pm. For the mPFC-DS group, C21 was delivered at 3:30 and 9:30pm, while SalB was delivered at 3:30 and 6:30pm. Drug was delivered on day 2 and/or 4 of ABA2 (Fig. 1B). Drugs were also delivered on post-recovery days, to test the effects of cell activation/suppression in the context of *ad libitum* food availability (Fig. 1B). Additionally, either C21 or SalB was delivered ~2hrs before perfusion, so as to reveal the extent of drug-induced cell activity/suppression by immunohistochemical detection of c-Fos protein within the virally infected portion of mPFC.

#### Statistical analyses

For data assessing two experimental groups (CON vs CaMKII or CON vs mPFC-DS), an F test was performed to establish whether standard deviations (SDs) were significantly different between groups. If SDs were not significantly different (p>0.05), an unpaired, two-tailed t-test was performed. If SDs were significantly different (p<0.05), an unpaired, two-tailed t-test with Welch’s correction was performed. Statistical analysis was performed and graphs were produced using the software Prism v. 6 or v. 8 (GraphPad Software, Inc.). Correlation data was assessed by simple linear regression, and Pearson R values and P values were reported.

## RESULTS

### ABA induction of CON mice reveals daily increase in FAA

As described in the methods, three different subgroups were used for CON mice: ‘GFP-CON’ mice (N=5; injected with CaMKII-GFP virus into the mPFC), ‘No-cre-CON’ (N=6; viral injection of cre-dependent DREADDs but no cre virus), and ‘No-Drug-cre-CON’ (N=5; viral injection of cre-eGFP virus and no DREADD or drug delivery). There were no significant differences between any of the CON subgroups, either in running distance or duration, during FAA (4pm-7pm) or food availability (7-9pm).

During ABA1, CON mice lost body weight significantly as compared to baseline, defined as the measure taken immediately prior to the start of ABA1 at 1pm., and mice were unable to regain body weight during the 2 hours of food availability (Fig. 1D). During ABA2, CON mice likewise lost body weight significantly due to restricted food access. Unlike during ABA1, mice during ABA2 were able to regain body weight to baseline levels by the end of the 2 hours of food access of FR day1 and 2 (FR1, 9 pm; FR2, 9 pm in Fig. 1D), but not during FR3 or FR4.

During ABA1, CON mice significantly increased their hourly wheel running distance, averaged over the course of the day, with a 41.5% increase above baseline during FR1 (t(30) = 2.414, p = 0.022, unpaired t-test) and a 58.0% increase above baseline during FR2 (t(30) = 2.614, p = 0.014, unpaired t-test), but which subsequently declined to a level no longer different from baseline by FR3 (t(30) = 1.463, p = 0.154, unpaired t-test; Fig. 1E). By contrast, hourly running distance averaged over the course of the three hours of FAA (4pm-7pm) increased steadily throughout ABA1. Because FR is imposed at 1pm, the 4pm-7pm FAA on FR1 occurred after only 3 hours of daylight food restriction. As expected, this extent of FR did not evoke any significant change in behavior. However, by FR2, the average hourly FAA running had increased by 341% above FR1 (t(18.2) = 2.360, p = 0.030, Welch’s t-test; Fig. 1E), and by FR3, percent increase was up to 584% above FR1 (t(16.1) = 2.923, p = 0.010, Welch’s t-test).

During ABA2, hourly running distance of CON mice, averaged over the course of the day, did not increase, unlike what was observed during ABA1. Instead, there was a significant decrease of 27.7% on FR2 (t(28) = 2.40, p = 0.023, unpaired t-test) and 27.3% on FR4 (t(30) = 2.73, p = 0.011, unpaired t-test), vs baseline (Fig. 1E). By contrast, hourly running distance averaged over the course of FAA, increased robustly throughout ABA2 – by 1258% on FR3 (t(15.46) = 2.864, p = 0.012, Welch’s t-test) and by 1505% on FR4 (t(15.47) = 3.452, p = 0.003, Welch’s t-test), vs FR1. Total running over the four days of ABA2 correlated positively with percent weight loss over the same timeframe (p = 0.010, R = 0.661, Fig. 1F), indicating that high runners were most at risk for precipitous weight loss. Note that FR3 and FR4 were the days when hourly FAA exceeded the hourly running averaged over the course of the day. On these days, the average daily FAA running correlated positively with daily percent weight loss (p = 0.014, R, = 0.602, Fig. 1G). By contrast, daily post-prandial running (9pm-1am, Fig. 1C) over FR3-4 did not correlate with daily percent weight loss (p = 0.361, R = 0.245).

The increase in FAA over the course of FR is not a novel finding (Chowdhury TG *et al.* 2013), but the role of PFC pyramidal neurons in FAA remained unknown. To test the hypothesis that these pyramidal neurons evoke FAA, we used C21 to activate Gq-DREADD expressed by PFC pyramidal neurons on FR2, when FAA running is relatively lower, and suppressed cell firing on FR4 using SalB KORD-DREADD, when FAA running is relatively higher, so as to minimize ceiling and floor effects that could obscure DREADD ligand effects.

### C21/Gq-DREADD activation of the subset of mPFC pyramidal cells projecting to DS increases FR-evoked running during FAA

For the mPFC-DS group, delivery of C21 during FAA of FR2 (4pm to 7 pm) significantly increased running distance by 245% vs CON subjects (t(19) = 3.085, p = 0.006, unpaired t-test), with significant between-group differences during the hour bins of 4-5pm (t(19) = 2.117, p = 0.048, unpaired t-test) and 5-6pm (t(19) = 4.006, p < 0.001, unpaired t-test). Running duration over the 4-7pm FAA period also increased by 194% (t(19) = 3.206, p = 0.010, unpaired t-test), compared to CON subjects (Fig. 2A). By contrast, delivery of C21 to the same mice during the same temporal period but after body weight restoration had no effect on running distance (t(14) = 0.705, p = 0.492, unpaired t-test) or running duration (t(14) = 0.202, p = 0.843, unpaired t-test; Fig. 2B). Thus, C21 delivery specifically increases FAA running during FR, but has no effect on running in general, when food is freely available. DREADD-driven effects observed during FAA of FR2 were no longer present during FAA of FR3, 18 hrs after the last ligand delivery (vs CON; for distance: t(23) = 1.208, p = 0.239, unpaired t-test; for duration: t(23) = 1.097, p = 0.284, unpaired t-test).

**Figure 2.**
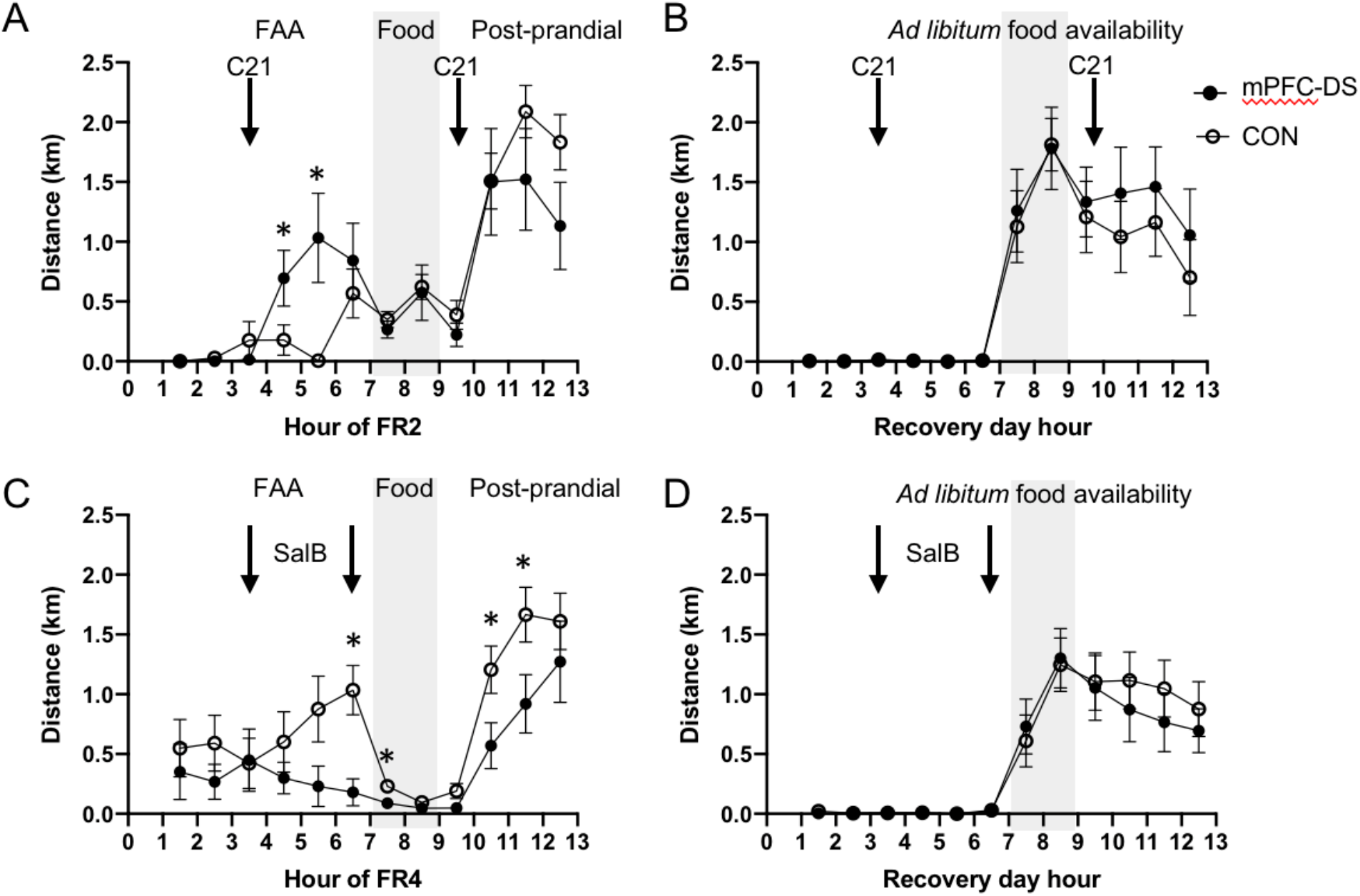
Bi-directional mPFC-DS control of FAA running occurs only during food restriction, not following recovery. C21 was administered to drive activity in mPFC-DS cells on FR2 **(A)** and following weight recovery **(B)**. SalB was administered to suppress activity in mPFC-DS cells on FR4 (**C**) and following weight recovery **(D)**. Injection times are indicated by black arrows. Hours of the day are indicated in the X-axis, with the 7-9pm hours of food availability highlighted in grey. Error bars indicate mean ± SEM; asterisks indicate significant differences, compared to CON values (p<0.05).

As previously reported (Chen YW, H Actor-Engel, *et al.* 2018), running distance and duration decreased dramatically for all animals following transition from FAA hours to feeding time (7pm-9pm). Delivery of C21 on FR2 had no further effect on feeding time (7pm – 9pm) running distance (t(19) = 0.508, p = 0.617, unpaired t-test) or duration (t(7.694) = 0.540, p = 0.604, Welch’s t-test; Fig. 2A), compared to CON subjects. Likewise, delivery of C21 to the same animals during the same hours but after weight restoration had no effect on running distance (t(14) = 0.133, p = 0.896, unpaired t-test) or running duration (t(14) = 0.126, p = 0.901, unpaired t-test; Fig. 2B).

Delivery of C21 on FR2 had no effect on post-prandial (9pm – 1am) running distance (t(19) =1.170, p = 0.257, unpaired t-test) or duration (t(7.321) = 1.181, p = 0.274, Welch’s t-test; Fig. 2A), relative to CON subjects’ post-prandial running. Likewise, delivery of C21 to the same animals after weight restoration had no effect on 9pm-1am running distance (t(14) = 0.747, p = 0.467, unpaired t-test) or running duration (t(14) = 0.529, p = 0.605, unpaired t-test; Fig. 2B). Thus, selective activation of DS-projecting mPFC cells increases FAA running without affecting subsequent running during the two hours of food availability or the post-prandial period.

### SalB/KORD suppression of the subset of mPFC pyramidal cells projecting to DS reduces FR-evoked running

Delivery of SalB on FR4 significantly decreased running distance during FAA (4pm to 7 pm) by 71.6% (t(20.04) = 2.427, p = 0.025, Welch’s t-test; Fig. 2C), with a significant decrease at the 6-7pm time bin immediately preceding food availability (t(23) = 2.944, p = 0.007, unpaired t-test). Likewise, running duration decreased by 58.5% (t(22.15) = 2.204, p = 0.038, Welch’s t-test). By contrast, delivery of SalB in recovered mice during the same hours had no effect on running distance (t(17) = 0.124, p = 0.231, unpaired t-test) or duration (t(17) = 0.697, p = 0.500, unpaired t-test; 2D), compared to CONs’. Thus, SalB delivery specifically decreases FAA running in the context of FR, but has no effect on running in general during *ad libitum* food availability.

Delivery of SalB on FR4 significantly decreased running distance during the feeding time (7pm to 9pm) by 59.3% (t(23) = 2.225, p = 0.036, unpaired t-test), with a significant decrease during the first hour of food availability (7-8pm time bin; (t(23) = 2.083, p = 0.048, unpaired t-test), but no effect on running duration t(23) = 0.983, p = 0.336, unpaired t-test; Fig. 2C). Delivery of SalB in weight-restored recovered mice during the same hours had no effect on running distance (t(17) = 0.161, p = 0.987, unpaired t-test) or duration (t(17) = 0.055, p = 0.956, unpaired t-test; Fig. 2D).

Delivery of SalB on FR4 significantly decreased post-prandial (9pm – 1am) running distance by 39.8%, compared to CON (t(23) =2.188, p = 0.039, unpaired t-test, Fig. 2C), with a significant decrease during the hours of 10-11pm (t(23) =2.102, p = 0.047, unpaired t-test) and 11pm-12am (t(23) =2.087, p = 0.048, unpaired t-test). SalB delivery also decreased post-prandial running duration by 36.3% (t(23) = 2.348, p = 0.028, unpaired t-test;). By contrast, delivery of SalB to the same animals during the same hours but after weight restoration had no effect on running distance (t(17) = 1.460, p = 0.2678, unpaired t-test) or running duration (t(17) = 0.987, p = 0.337, unpaired t-test; Fig. 2D). Thus, SalB delivery decreases running during FAA, food allowance, and post-prandial periods, specifically during FR, without affecting running during recovery.

### Bidirectional modulation of the mPFC-DS pathway does not alter body weight or food consumption

In order to determine the impact of wheel running upon body weight, weight loss was calculated as the percent change in weight between the baseline weight measured immediately before the start of FR1 and the FR weight measured at 7pm, immediately following FAA. DREADD activation of mPFC-DS cells via C21 on FR2 did not measurably alter the percent weight loss from baseline (t(23)=0.468; p=.644, unpaired t-test) or alter percent weight gain over the two hours of food allowance (t(23)=0.150; p=.882, unpaired t-test). Likewise, suppression of mPFC-DS cells on FR4 did not measurably alter percent weight loss from baseline (t(23)=0.023; p=.982, unpaired t-test) or alter percent weight gain over the two hours of food allowance (t(23)=0.549; p=.589, unpaired t-test). Food consumption over the period of food allowance also was not different between mPFC-DS and CON groups following cell activation on FR2 (t(10.33)=1.357; p=.204, Welch’s t-test) or suppression on FR4 (t(23)=1.159; p=.258, unpaired t-test).

### C21 activation of mPFC pyramidal cells via CaMKIIα promoter-driven Gq-DREADD increases FR-evoked running during FAA

We then tested whether activation of the full population of mPFC pyramidal cells would increase FAA hyperactivity, or broaden the hyperactive response to include running during *ad libitum* food availability. After delivery of C21during FAA of FR2 to activate all mPFC pyramidal cells targeted by the CaMKIIα promoter, we observed a 255% increase in running distance (t(20) = 2.919, p = 0.009, unpaired t-test) vs CON subjects, with a significant increase at the 5-6pm time bin (t(20) = 3.744, p = 0.001, unpaired t-test). Running duration increased by 190% (t(20) = 3.152, p = 0.005, unpaired t-test), compared to CON subjects’ (Fig. 3A). DREADD-driven effects observed on FR2 were no longer present on FR3, 24 hours after C21 administration (for distance: t(22) = 0.631, p = 0.534, unpaired t-test; for duration: t(22) = 0.821, p = 0.421, unpaired t-test).

**Figure 3.**
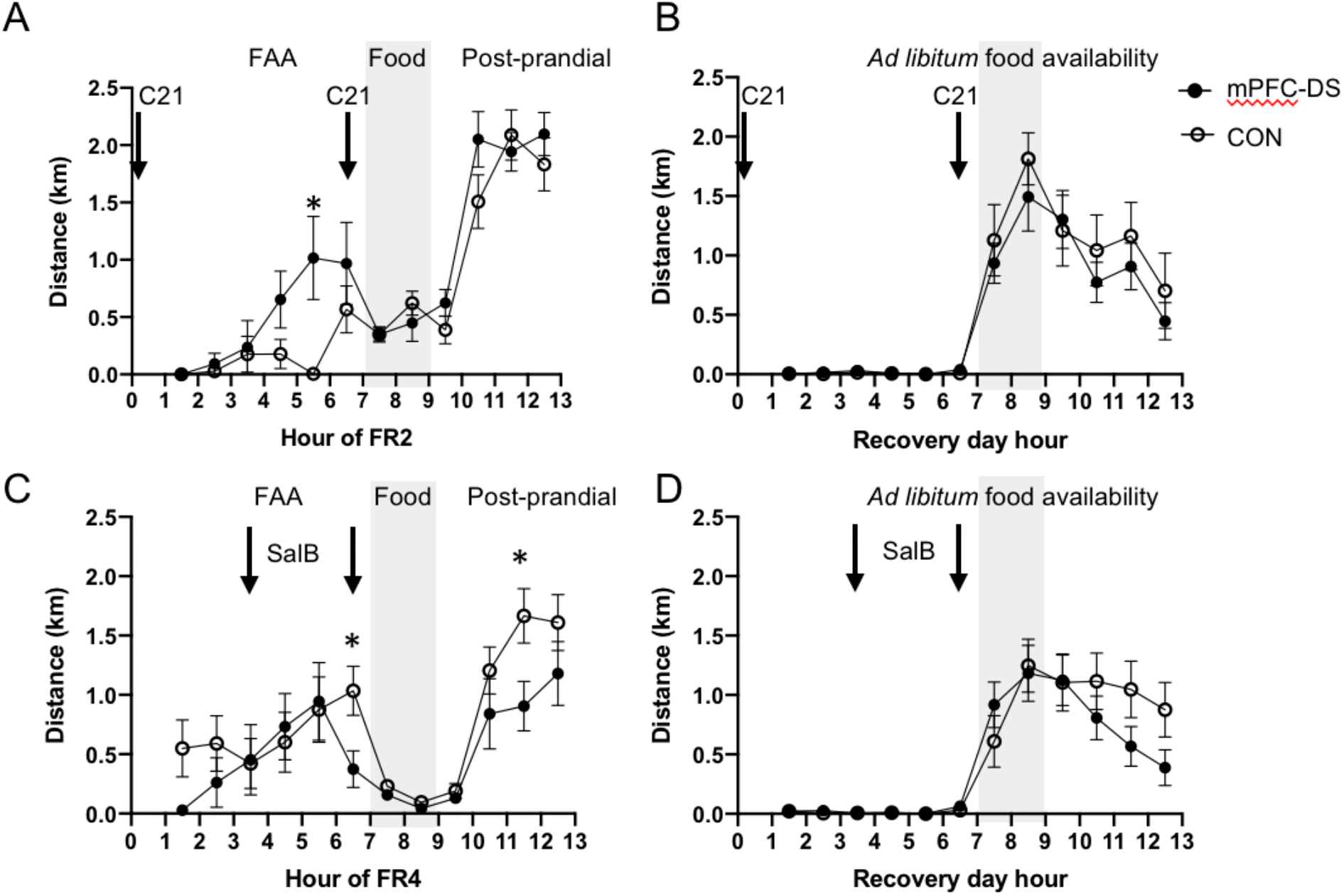
Activation of general mPFC pyramidal cells drives FAA running during food restriction, but not following recovery, while suppression has a broader temporal effect. C21 was administered to drive activity of mPFC pyramidal cells on FR2 (**A)** and following weight recovery **(B)**. SalB was administered to suppress activity of mPFC pyramidal cells on FR4 (**C**) and following weight recovery (**D**). Injection times are indicated by black arrows. Hours of the day are indicated in the X-axis, with the 7-9pm hours of food availability highlighted in grey. The values are mean ± SEM. Asterisks indicate significant differences, compared to CON values (p<0.05).

The C21 effect on running distance was specific to the FR state, as delivery of C21 to the same animals after weight restoration had no effect on running distance during the same 4-7pm hours of FAA (t(20) = 0.750, p = 0.462, unpaired t-test, Fig. 3B). Running duration from 4-7pm in recovered CaMKII subjects treated with C21 differed from CON subjects’ by 154% (t(17.99) = 2.952, p = 0.009, Welch’s t-test). However, the C21 effect on running duration during recovery was relatively small compared to the effect during FR, with a difference in means of only 6.587 ± 2.231 active minutes during recovery, vs 44.27 ± 14.05 active minutes during FR.

For the CaMKII group, delivery of C21 during FR2 food allowance (7 pm to 9 pm) had no effect on running distance (t(20) = 0.837, p = 0.412, unpaired t-test) or duration (t(20) = 1.332, p = 0.198, unpaired t-test), compared to CON subjects (Fig. 3A). Likewise, delivery of C21 in weight-restored recovered mice during the same hours had no effect on running distance (t(20) = 0.871, p = 0.394, unpaired t-test) or duration (t(20) = 0.896, p = 0.381, unpaired t-test; Fig. 3B). Thus, activation of mPFC pyramidal cells via C21 delivery specifically increases FAA running without affecting subsequent running during the period of food allowance.

Delivery of C21 on FR2 had no effect on post-prandial (9pm – 1am) running distance (t(20) =0.994, p = 0.332, unpaired t-test), or duration, although there was a trend towards increase for the latter (t(20) = 1.994, p = 0.060, unpaired t-test; Fig. 3A). Likewise, delivery of C21 to the same animals after weight restoration had no effect on 9pm-1am running distance (t(20) = 0.592, p = 0.560, unpaired t-test) or running duration (t(20) = 0.330, p = 0.745, unpaired t-test; Fig. 3B). Thus, activation of CaMKII pyramidal cells specifically increases FAA running without affecting subsequent running during the two hours of food availability or the post-prandial period. Together, this indicates that the robust C21 effect observed for activation of mPFC-DS projecting cells was matched, but not enhanced, when the suppression included mPFC pyramidal cells that project elsewhere.

### SalB suppression of mPFC pyramidal cells via CaMKIIα promoter-driven KORD-DREADD decreases FR-evoked running

For the CaMKII group, delivery of SalB on FR4 had no effect on pooled FAA (4-7pm) running distance (t(24) = 0.452, p = 0.655, unpaired t-test) or duration (t(24) = 0.367, p = 0.717, unpaired t-test), compared to CON subjects’ (Fig. 3C). However, there was a significant 40% decrease in running distance specifically at the 6-7pm time bin, when hunger is most intense (t(24) = 2.272, p = 0.032, unpaired t-test). Delivery of SalB to mice after weight restoration had no effect on running distance (t(10.12) = 1.201, p = 0.257, Welch’s t-test) or duration (t(17) = .1.049, p = 0.309, unpaired t-test; Fig. 3D).

For the CaMKII group, delivery of SalB during food allowance of FR4 (7 pm to 9 pm) had no effect on running distance (t(24) = 1.645, p = 0.113, unpaired t-test) or duration (t(24) = 0.907, p = 0.374, unpaired t-test), when compared to CON subjects’ (Fig. 3C). Likewise, delivery of SalB in recovered mice during the same hours produced no effect on running distance (t(24) = 0.108, p = 0.915, unpaired t-test) or duration (t(24) = 0.597, p = 0.644, unpaired t-test; Fig. 3D).

Following delivery of SalB on FR4, CaMKII mice trended towards a reduction of post-prandial (9pm – 1am) running distance (t(24) =1.796, p = 0.085, unpaired t-test), with a significant decrease occurring for the hour of 11pm-12am (t(24) =2.267, p = 0.033, unpaired t-test). There was no significant reduction in post-prandial running duration (t(24) = 1.427, p = 0.167, unpaired t-test; Fig. 3C). Likewise, delivery of SalB during the same time bin but after weight restoration had no significant effect, but a strong trend toward reduction for post-prandial running distance (t(24) = 1.943, p = 0.064, unpaired t-test) and running duration (t(24) = 1.905, p = 0.069, unpaired t-test; Fig. 3D). Thus, suppression of mPFC pyramidal cells via SalB delivery produces a measurable reduction in wheel running only during the last hour of FAA, when hunger is most intense, and during the 11-12 hour post-prandial time bin. This indicates that the SalB effect upon mPFC pyramidal cells of reducing FAA was present, but much muted, when the suppression included mPFC pyramidal cells that project elsewhere.

### Bi-directional modulations of mPFC pyramidal cells via CaMKIIα promoter-driven Gq-DREADD and KORD-DREADD do not alter body weight or food consumption

Up to the time of entry into ABA2, CON and chemogenetically modulated groups of mice exhibited no group difference in body weight or food consumption. Specifically, there was no significant differences in weight on ~P36 when mice were introduced to wheels, between mPFC-DS vs CON mice (t(22) = 1.04, p = 0.310, unpaired t-test) or CaMKII vs CON mice (t(25) = 0.155, p = 0.878, unpaired t-test). Likewise, there was no significant group differences in weight (mPFC-DS vs CON: t(22) = 1.316, p = 0.201, unpaired t-test; CaMKII vs CON: t(27) = 0.780, p = 0.442, unpaired t-test) or running distance (mPFC-DS vs CON: t(17) = 0.069, p = 0.946, unpaired t-test; (CaMKII vs CON: t(24) = 1.057, p = 0.301, unpaired t-test) at termination of ABA1, indicating no major differences in vulnerability to consider between groups as they entered ABA2 for chemogenetic manipulations.

Despite the robust effect that DREADD activation of mPFC pyramidal cells via C21 during ABA2 had on FAA, this change in behavior did not measurably alter percent weight loss for FR2 (t(7.54)=0.206; p=.846, Welch’s t-test) or alter percent weight gain over the two hours of food allowance (t(22)=0.404; p=.690, unpaired t-test), relative to CON subjects. Likewise, suppression of mPFC pyramidal cells by SalB on FR4 did not measurably alter percent weight loss on FR4 (t(24)=1.639; p=.114, unpaired t-test), or percent weight gain over the two hours of food allowance on FR4 (t(24)=.147; p=.885, unpaired t-test), compared to CON. Food consumption during the two hour window also was not different between CaMKII and CON groups following cell activation on FR2 (t(9.016)=.636; p=.540, Welch’s t-test) or suppression on FR4 (t(24)=1.367; p=.184, unpaired t-test).

### Elevated plus maze

Elevated plus maze was administered to all subjects to assess general anxiety level before either DREADD ligand was administered, during the hours of 6am-9am of FR1. As compared to CON subjects, there were no significant differences in percent duration of time spent in the open arms of the EPM for either the CaMKII (t(29)=0.688; p=.497, unpaired t-test) or mPFC-DS groups (t(10.42)=1.118; p=.289, Welch’s t-test), thus confirming that the effect observed following DREADD ligand administration had no contribution from pre-existing differences in the general anxiety level.

### Viral expression

Virally mediated expression of cre-dependent DREADDs was observable in layers 2-6 of mPFC (Fig. 4A-B). The mCherry reporter identifies expression of the Gq receptor, and the mCitrine reporter identifies expression of the KORD receptor. Of all 18 hemispheres of the mPFC-DS group, 94% expressed both viruses in the Cg1, 100% expressed both viruses in the PL, and 94% expressed both viruses in the IL. Cre-dependence of the viral expression was confirmed, as cre-dependent DREADD virus was unable to replicate in the absence of the retrogradely transported (rg)-cre-virus (N=5; Fig. 4G). We confirmed mCitrine positive (KORD receptor-expressing) axon terminals in the grey matter of the medial DS (Fig. 4H). Several of the cells supported viral expression of both mCherry and mCitrine, indicating that a single cell can support replication of both cre-dependent DREADD viruses as well as the rg-cre virus without visible damage to cells (Fig. 4B). The absence of DREADDs in GABA-INs was confirmed by the absence of GAD/DREADD co-labeling (Fig. 4C).

**Figure 4.**
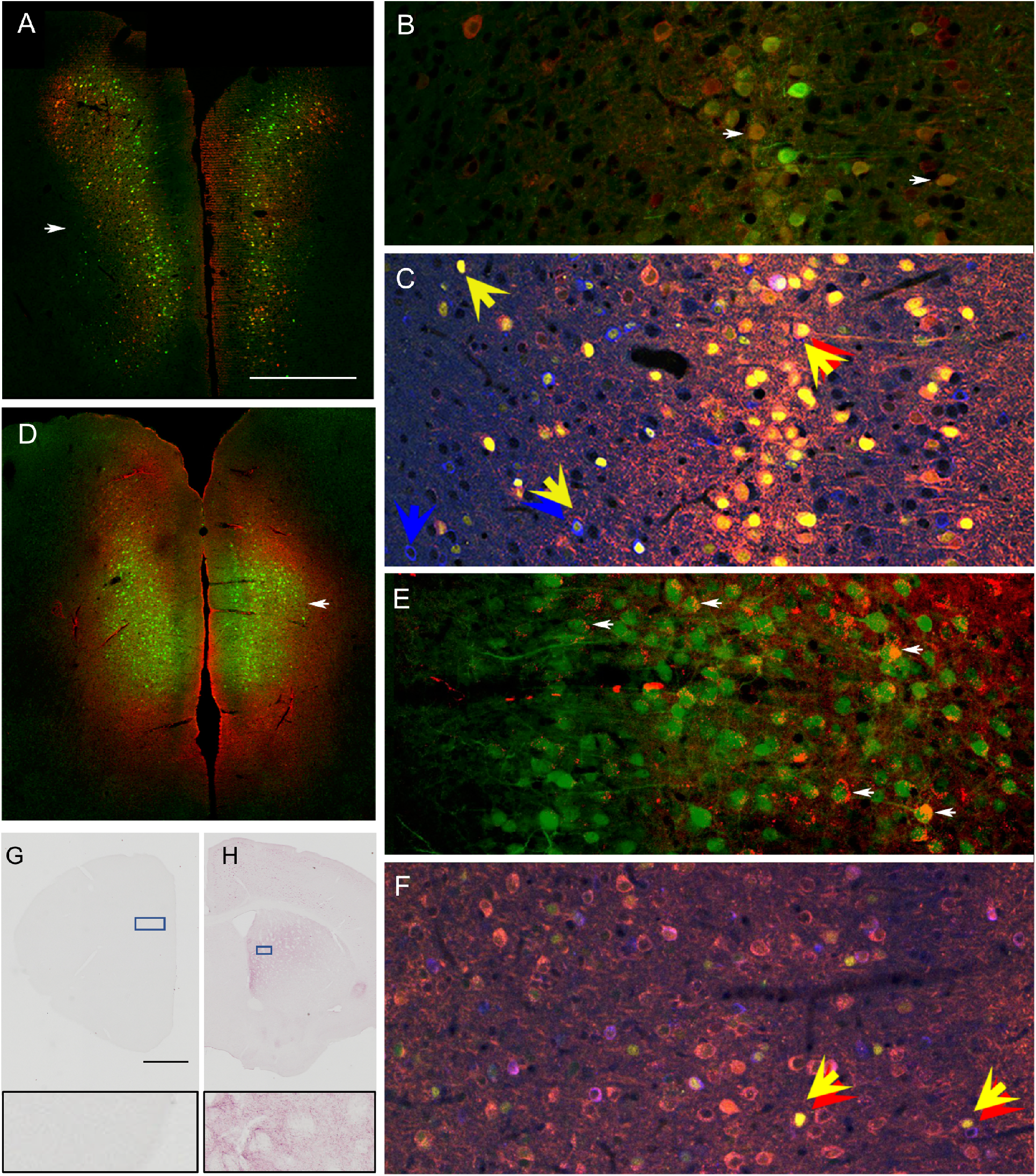
Immunofluorescence reveals pyramidal cell-specific DREADD expression and verifies DREADD-driven modulation of pyramidal cell activity. mCherry and mCitrine protein immunoreactivity in the mPFC reflect the virally mediated transfer of genes encoding DREADDs that were either excitatory (Gq-mCherry) or inhibitory (KORD-mCitrine). Viral expression was either Cre-dependent with retrograde Cre delivered into the DS, for expression specifically in pyramidal neurons projecting to DS (mPFC-DS; panels A and B) or CaMKII-promoter driven (panels D and E) for pyramidal neuron specific expression. Panels B and E show details of the prelimbic (PL) subregion of mPFC (arrows in panels A and D). mCherry-immunoreactivity (red) appears as puncta in the perikaryal cytoplasm. In contrast, mCitrine-immunoreactivity (green) appears to fill the nucleus, perikaryal cytoplasm and apical dendrites of pyramidal neurons. Arrows in B and E point to some of the dually immunolabeled cell bodies. **C and F**. Triple immunofluorescence was performed to simultaneously detect c-Fos protein in nuclei (yellow, examples highlighted by yellow arrows) and mCherry (red, red arrows), together with glutamic acid decarboxylase (GAD), the rate-limiting synthetic enzyme of GABAergic interneurons (GABA-INs, blue, blue arrows) in the PL by confocal microscopy. Ante-mortem C21 was delivered to activate DREADD-Gq receptors 2-3 hours prior to euthanasia. mCitrine, the KORD virus reporter (green), was also detected, but not amplified by IF. **Panel C** depicts PL of mPFC-DS tissue. **Panel F** depicts PL of tissue transfected with the CaMKIIα-Gq-DREADD-mCherry transgene. Overall, c-Fos immunoreactivity is more prevalent following activation of the mPFC-DS subpopulation, relative to activation of the general population of pyramidal neurons with CaMKIIα-Gq-DREADD-mCherry. GABA-INs with c-Fos in nucleus are also apparent (yellow-blue double arrow), suggesting recruitment of GABA-INs. **G**. Absence of KORD viral replication in mCitrine-expressing cells in mPFC of ‘No-cre-CON’ after immunohistochemical detection of mCitrine, the reporter for KORD, using an anti-GFP antibody. The rectangle at the bottom of panel E shows mPFC at a higher magnification. **H.** mCitrine-expressing axons in dorsal striatum, revealed by immunohistochemical detection of mCitrine, using anti-GFP, and visualized by the HRP reaction product, VIP from a section of a brain transfected with KORD-mCitrine. Anti-GFP also detected the eBFP reporter for the retrograde cre virus, localized to nuclei of transfected pyramidal neurons. The insets of G and H correspond to the boxes in the panels. Calibration bar equals 100 μm in panels B, C, E, and F and 600 μm in panels A and D. Calibration bar in panel G applies to both panels G and H, equaling 100 μm.

Virally mediated expression of CaMKIIα promoter-driven DREADDs was observable in layers 2-6 of mPFC (Fig. 4D-E). Viral expression spanned the dorsal-ventral axis of the mPFC, centering around the prelimbic cortex (PL) subregion, with anteroposterior axis center of 2.34+ Bregma. Of all 30 hemispheres targeted in the CAMKII group, 100% expressed both viruses in the cingulate cortex (Cg1) and PL regions, and 96% expressed both viruses in the infralimbic cortex (IL). Virally mediated expression of CaMKIIα promoter-driven DREADDs was found specifically in pyramidal cells, identifiable by their distinct cell body shape and singular apical dendrites. Several of the cells supported viral expression of both mCherry and mCitrine, indicating that a single cell could support replication of both viruses without visible detriment to cell health (Fig. 4E).

### Expression of c-Fos after DREADD ligand delivery

Immunocytochemistry was conducted to assess the extent to which subcutaneous drug delivery 2 hours prior to euthanasia could alter c-Fos activity in DREADD-expressing cells, as well as GABA-INs (Fig. 4C and F). Basal c-Fos activity in mPFC pyramidal cells was assessed in the GFP-CON mice (expressing GFP but not DREADD in mPFC pyramidal cells; N=5), which received C21 two hours before euthanasia. For these subjects, 7.7% ± 1.6% of GFP+ pyramidal cells expressed c-Fos.

For the CaMKII subjects that received SalB prior to perfusion (CaMKII-SalB; N=7), c-Fos immunoreactivity was evident within 3.9% ± 2.6% of pyramidal neurons, identified by GFP immunoreactivity to the KORD-reporter protein, mCitrine. This level of c-Fos immunoreactivity did not differ significantly from basal mPFC c-Fos expression (t(10) = 1.153, p = 0.276, unpaired t-test), although we did detect a floor effect, as 4 out of the 7 CaMKII-SalB cases had zero c-Fos expression among KORD/mCitrine/GFP expressing cells, while none of the 5 GFP-CON animals exhibited zero values. Of the 9 mPFC-DS group mice, only two received SalB prior to perfusion, with c-Fos immunoreactivity evident within 2.8% ± 1.0% of mCitrine/GFP+ cells.

Despite the lack of C21-driven running in weight recovered mice with *ad libitum* food access, we did observe a C21-driven increase in pyramidal cell c-Fos expression in weight-recovered mice for both experimental groups. Of the CaMKII subjects that received C21 prior to perfusion (CaMKII-C21; N=8), c-Fos expression in pyramidal cells was 2.55 times greater vs. GFP-CON, with 19.7% ± 4.1% of mCherry positive cells expressing c-Fos (t(8.898) = 2.731, p = 0.023, Welch’s t-test, Fig 5A). This ratio is consistent with previously reported data using CaMKII-DREADD to activate pyramidal cells in the mPFC (Pati S et al. 2018). By contrast, pyramidal c-Fos expression was 10.67 times greater in the C21-activated mPFC-DS group (N=7), with 82.5% ± 5.2% of mCherry positive cells expressing c-Fos (t(7.055) = 13.63, p < 0.001, Welch’s t-test, Fig 5B).

**Figure 5.**
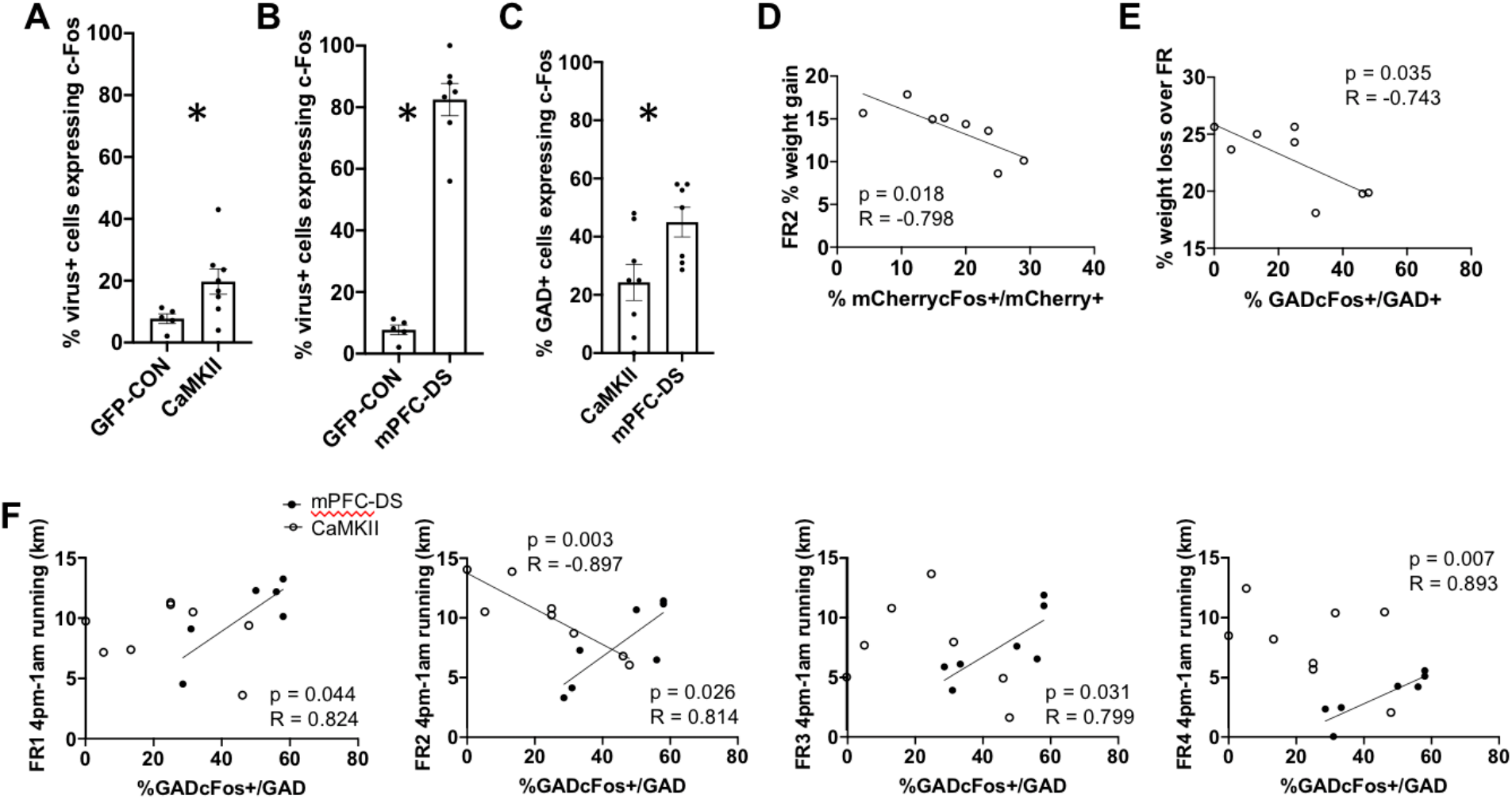
Quantification of c-Fos, GAD and mCherry reveals microcircuitry activated by C21 and correlation of microcircuitry activity with behavior. **A.** C21-driven c-Fos expression is greater in the pyramidal cells of the CaMKIIα-Gq-DREADD group, vs the CaMKIIα-GFP-control group (GFP-CON). **B.** C21 driven c-Fos expression is greater in the pyramidal cells of the mPFC-DS group, vs the CaMKIIα-GFP-control group. **C.** C21-driven recruitment of GABA-INs is greater for the mPFC-DS group vs the CaMKII group. Asterisks in panels D, E and F denote p<0.05. Values of the graphs are mean ± SEM. **D**. In the CaMKII group, percent of mCherry+ pyramidal cells expressing c-Fos correlates negatively with percent weight gain on FR2, but not other days. **E.** In the CaMKII group, percent of GAD+ GABA-INs expressing c-Fos correlated negatively with weight loss over the four-day course of FR. **F**. In the CaMKII group (open circle), percent of GABA-INs expressing c-Fos correlates significantly and positively with running that spans 4pm-1am (includes FAA, food availability, and post-prandial time bins) during C21 activation on FR2. In the mPFC-DS group (filled circle), the percent of the GABA-INs expressing c-Fos correlates significantly and positively with running that spans 4pm-1am on all 4 days of FR. Each graph shows the outcome of the correlation analysis for each of the four days of food restriction. Solid line indicates significant correlation, no line indicates p > 0.1.

For explanation of the difference in c-Fos immunoreactivity of the mPFC-DS group versus the CaMKII group, we turned to mPFC microcircuitry, and asked whether C21 delivery evoked a different response when acting on the mPFC-DS subpopulation, vs general pyramidal cell activation. Interestingly, the percent of GAD-immunoreactive cells co-expressing c-Fos was greater following antemortem C21 delivery to the mPFC-DS group (45.0%), vs. the CaMKII group (24.3%; (t(13) = 2.529, p = 0.025, unpaired t-test; Fig. 5C), indicating that, compared to the general pyramidal cell population, activity from the mPFC-DS subpopulation evokes a greater GABA-IN response.

### Individual differences in neuronal activity as revealed by c-Fos expression relate to individual differences in wheel running

While C21 evoked robust 245% and 255% increase in FR2 wheel running via activation of the mPFC-DS and the general population of mPFC pyramidal cells, respectively (Figs 2 and 3), the same measurements also revealed multi-fold differences in individuals’ extent of C21-evoked running. To better understand the nature of these individual differences, we assessed whether these differences related to the extent of C21/DREADD-driven c-Fos activity of neurons in the mPFC (see Table 1 for p and R values). Contrary to our expectation, c-Fos expression in mCherry+ pyramidal cells of either mPFC-DS or CaMKII groups did not correlate significantly with running distance during FAA or the 4pm-1am period. This indicates that another group of cells might contribute more strongly towards individual differences in running.

**Table 1.** Correlations between c-Fos expression and running/weight change. List of variables and the observed results of correlations for both the mPFC-DS subpopulation and general mPFC pyramidal cell population. Percent weight loss is calculated as the weight at 7pm on the FR day subtracted from the weight at the start of ABA2, all divided by the weight at the start of ABA2. Percent weight gain is calculated as the weight at 9pm subtracted from the weight at 7pm of the same day, all divided by the weight at 7pm.

| Variable | Variable |  | PFC-DS group |  | CaMKII group |  |
| --- | --- | --- | --- | --- | --- | --- |
|  |  |  | p value | r value | p value | r value |
| Percent of mCherry+ pyramidal cells expressing c-Fos | Running (km) | FR 1 | 0.665 | -0.228 | 0.967 | -0.180 |
|  |  | FR 2 | 0.374 | -0.400 | 0.096 | -0.627 |
|  |  | FR 3 | 0.099 | -0.671 | 0.673 | -0.178 |
|  |  | FR 4 | 0.325 | -0.439 | 0.582 | -0.231 |
|  | % weight loss | FR 1 | 0.068 | 0.720 | 0.861 | -0.074 |
|  |  | FR 2 | 0.774 | -0.134 | 0.449 | 0.314 |
|  |  | FR 3 | 0.579 | -0.256 | 0.096 | -0.628 |
|  |  | FR 4 | 0.587 | -0.251 | 0.186 | -0.521 |
|  | % weight gain over 2 hrs of food availability | FR 1 | 0.366 | 0.406 | 0.383 | 0.359 |
|  |  | FR 2 | 0.766 | -0.139 | **0.018** | **-0.798** |
|  |  | FR 3 | 0.539 | -0.283 | 0.741 | 0.140 |
|  |  | FR 4 | 0.225 | 0.277 | 0.464 | 0.304 |
| % GABA-IN expressing c-Fos | Running (km) | FR 1 | **0.044** | **0.824** | 0.654 | -0.189 |
|  |  | FR 2 | **0.026** | **0.814** | **0.002** | **-0.897** |
|  |  | FR 3 | **0.031** | **0.799** | 0.497 | -0.283 |
|  |  | FR 4 | **0.007** | **0.893** | 0.297 | -0.422 |
|  | % weight loss | FR 1 | 0.641 | -0.217 | 0.558 | -0.246 |
|  |  | FR 2 | 0.874 | 0.075 | 0.507 | 0.278 |
|  |  | FR 3 | 0.752 | 0.148 | **0.004** | **-0.883** |
|  |  | FR 4 | 0.844 | -0.092 | **0.035** | **-0.743** |
|  | % weight<br>gain over<br>2hrs of food<br>availability | FR 1 | 0.359 | -0.412 | 0.465 | -0.304 |
|  |  | FR 2 | 0.654 | -0.208 | 0.477 | -0.296 |
|  |  | FR 3 | 0.086 | 0.691 | 0.365 | 0.371 |
|  |  | FR 4 | 0.886 | 0.067 | 0.238 | 0.471 |

For the CaMKII group, the percent of GAD+ cells expressing c-Fos correlated negatively with running distance during the 4pm-1am hours of C21 activation on FR2, (Fig. 5F; Table 1). with no correlation between GAD+/cFos+ cells and running during this period of other FR days. Within this window of C21 activation on FR2, GAD+ c-Fos immunoreactivity correlated specifically with running during the 4-7pm FAA period (p = .003; R= −0.893), and not with running during the 2 hrs of food availability (p = 0.955; R = −0.024) or the post-prandial period (p =0.131; R = 0.581). This negative correlation, only on FR2, indicates that the C21-induced recruitment of GABA-INs by the general population of pyramidal cells contributed towards individual differences in the dampening of hyperactivity, particularly during the FAA hours.

By contrast, for the mPFC-DS group, the percent of GAD+ cells expressing c-Fos correlated *positively* with running. Moreover, in contrast to the CaMKII group, GAD+/c-Fos+ cells correlated with running during 4pm-1am not only on FR2, but on all FR days (Fig. 5F; Table 1). This effect was specific to running in the context of FR, as there was no correlation between running and GAD+ cell c-Fos expression during the days with *ad libitum* food access preceding FR (preceding FR by 1 day: p =0.387, R =0.436; preceding by two days: p = 0.713, R =-0.172) or on days after weight-restoration (with C21 delivery: p = 0.163, R=0.591; with SalB delivery: p =0.458, R=0.339; with no drug: p =0.114, R=0.650). Since C21-activation of pyramidal neurons in the mPFC augments FR-evoked running, the positive correlation between running and GAD+/c-Fos+ cells, which dampen pyramidal neurons, was contrary to expectation (reconciled in Fig. 10 and Discussion).

### Individual differences in neuronal activity as revealed by c-Fos expression relate to individual differences in weight loss over FR and weight gain during the hours of food access

For the CaMKII group, individual differences in percent weight loss correlated negatively with percent of GAD+ cells expressing c-Fos on FR3 and FR4 (Fig 5D; Table 1), indicating that GABAergic inhibition, recruited by C21 activation of pyramidal cells, may have been protective against weight loss during the days with severest weight loss. By contrast, for the mPFC-DS group, individual differences in percent weight loss did not correlate with percent of GAD+ cells expressing c-Fos on either day.

As stated earlier, analysis of weight gain revealed no group differences across the C21-treated versus CON for the CaMKII group or the mPFC-DS group. Yet, for the CaMKII group, the percent of mCherry+ pyramidal cells expressing c-Fos correlated negatively with the percent weight gain over the 2 hr period of food availability on FR2, when C21 was active, but not on other FR days (Fig 5E; Table 1), indicating that pyramidal cell activation by C21 on FR2 may have hampered food consumption. By contrast, the percent of DS-projecting mCherry+ pyramidal neurons expressing c-Fos showed no correlation. This could reflect greater C21 activation of non-DS-projecting mPFC pyramidal cells with targets to a feeding center, such as the lateral hypothalamus, which regulates food consumption when activated (Cassidy RM and Q Tong 2017) and receives strong projection from mPFC (Gabbott PL et al. 2005).

### Electron microscopic verification of Gq-DREADD expression

For neurons to be responsive to DREADD ligands, DREADDs need to be expressed at the plasma membrane. Electron microscopy verified that immunoreactivity for mCherry, the reporter for cre-dependent Gq-DREADD expression, was localized to the plasma membrane of cell bodies containing nuclei with smooth contour, characteristic of pyramidal neurons (Fig. 6A; White E 1989). Cre-dependent DREADD expression was also evident along the plasma membrane of dendritic spines, also indicating expression in excitatory pyramidal cells (Fig. 6B). Synapses formed by axon terminals with mCherry-immunoreactivity exhibited thick postsynaptic densities (PSDs; Fig. 6C), indicated that these synapses are excitatory, as would be expected for glutamatergic synapses formed by pyramidal neurons (White E 1989). mCherry-immunoreactive pyramidal neurons targeted dendritic spines, indicating pyramidal-to-pyramidal synaptic targets (Fig. 6D). mCherry-immunoreactive pyramidal neurons also targeted dendritic shafts that were mCherry-immunonegative (Fig. 6E). Since GABA-INs are known to receive excitatory synaptic inputs directly onto their dendritic shafts and cell bodies (White E 1989), this relationship was indicative of pyramidal-to-GABA-IN synaptic targets, thus providing the cellular substrate for c-Fos expression in GAD+ cells (Fig. 5C).

**Figure 6.**
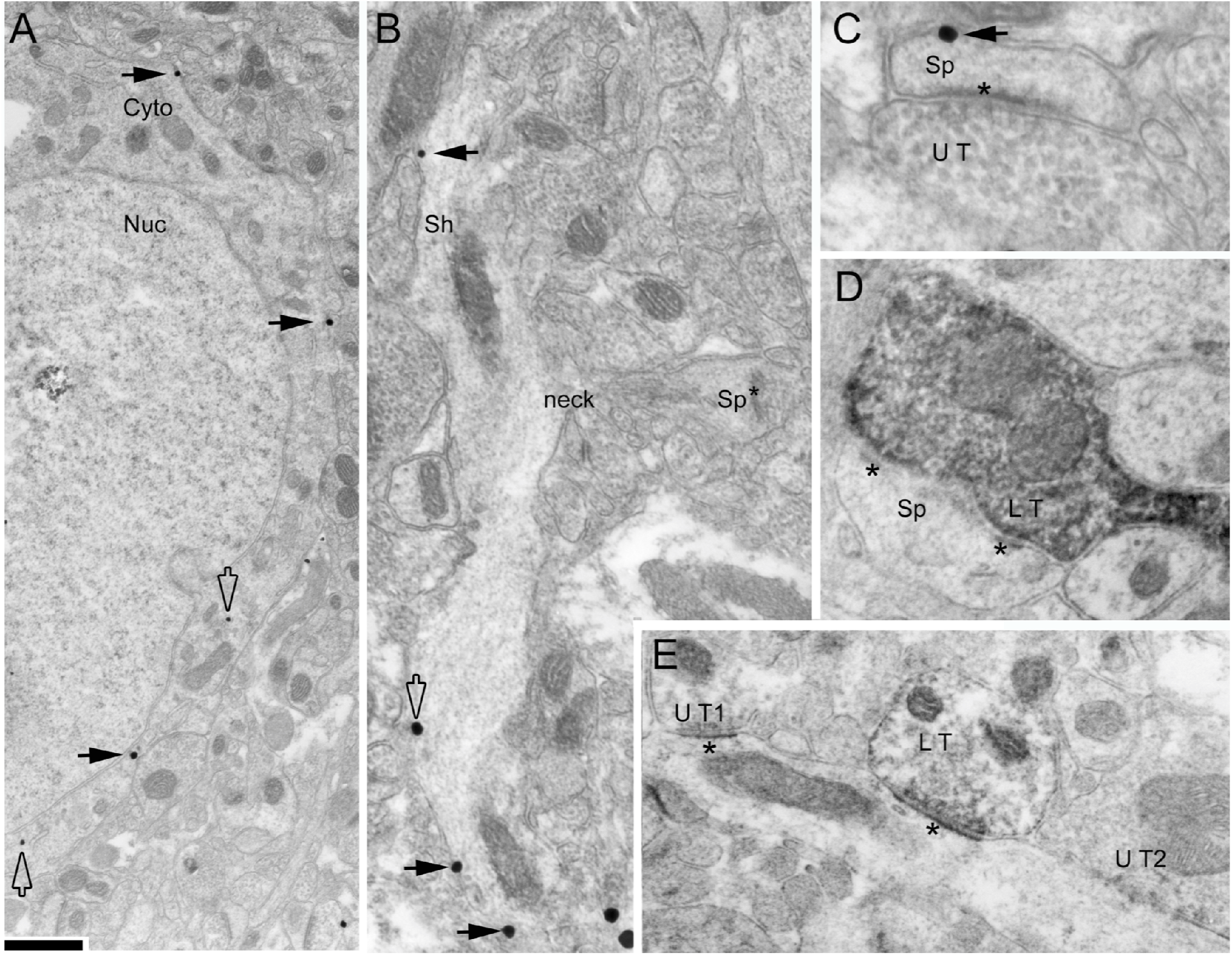
Electron microscopic verification of the localization of DREADD-mCherry to the plasma membrane. Silver-intensified gold labels (SIG; **Panels A through C**) and HRP-DAB reaction products (**Panels D and E**) were used to immunolabel mCherry, the reporter of the Gq-DREADD. The non-diffusible SIG reveals the location of mCherry along intracellular surfaces of the plasma membrane (black arrows in panel A and other panels) of a neuronal cell body, likely to be of a pyramidal neuron, based on the smooth contour of the nuclear (Nuc) envelope. The open arrows point to SIGs located in the cytoplasm (Cyto), removed from the plasma membrane. Plasmalemmal labeling for mCherry is evident in dendritic shafts (Sh, Panel B) from which a spine head forming an asymmetric synapse with thick postsynaptic density (PSD) protrudes via its spine neck, indicating that this dendrite is of a spiny neuron, presumably pyramidal. Spine heads (Sp, Panel C) forming asymmetric synapses with unlabeled terminals (UT) are immunoreactive for mCherry along the plasma membrane. mCherry also occurs presynaptically, in axon terminals forming asymmetric, putatively excitatory synapses with prominent postsynaptic densities (PSDs) onto dendritic spines (LT=labeled terminal, Panel D), indicative of a pyramidal-to-pyramidal synapse. Other mCherry axon terminals form asymmetric synapses onto dendritic shafts (Panel E), indicative of pyramidal-to-GABAergic interneuron synapse. Calibration bar = 1 μm for panel A, 480 nm in B, 240 nm in C, 300 nm in D, and 400 nm in E.

### Greater GABAergic innervation of non-DS projecting layer 5 mPFC pyramidal cells

Electron microscopy was then used in the mPFC-DS group (N=7), to assess GABAergic axo-somatic contacts to layer 5 mPFC pyramidal cells, either expressing mCherry, confirming projection to DS (Fig. 7), or lacking mCherry expression (Fig. 8). GABA-INs were identified by DAB precipitate, reflecting GAD immunoreactivity, while mCherry immunoreactivity was identified by another electron-dense immunolabel, SIG (silver-intensified colloidal gold). GABAergic innervation was quantified as percent of the somatic plasma membrane contacted directly by GAD+ terminals. Surprisingly, we found that the mCherry positive, DS-projecting subpopulation of mPFC pyramidal cells had 42.3% less axo-somatic GABAergic innervation than neighboring unlabeled pyramidal cells (t(12) = 2.914; p=0.013; unpaired t-test, Fig. 9A). This was due to lower density of GABAergic innervation (t(12) = 3.392; p=0.005; unpaired t-test, Fig. 9B), while size of synaptic terminal was not different between groups (t(12) = 0.243; p=0.812; unpaired t-test, Fig. 9C). Together with the c-Fos data on GABA-INs, this indicates that, although driving mPFC-DS pyramidal cells elicits a strong response from GABA-INs in the mPFC s, GABAergic action has stronger axo-somatic target to neighboring non-DS projecting mPFC pyramidal cells, vs self-inhibition (Figure 10 for summary).

**Figure 7.**
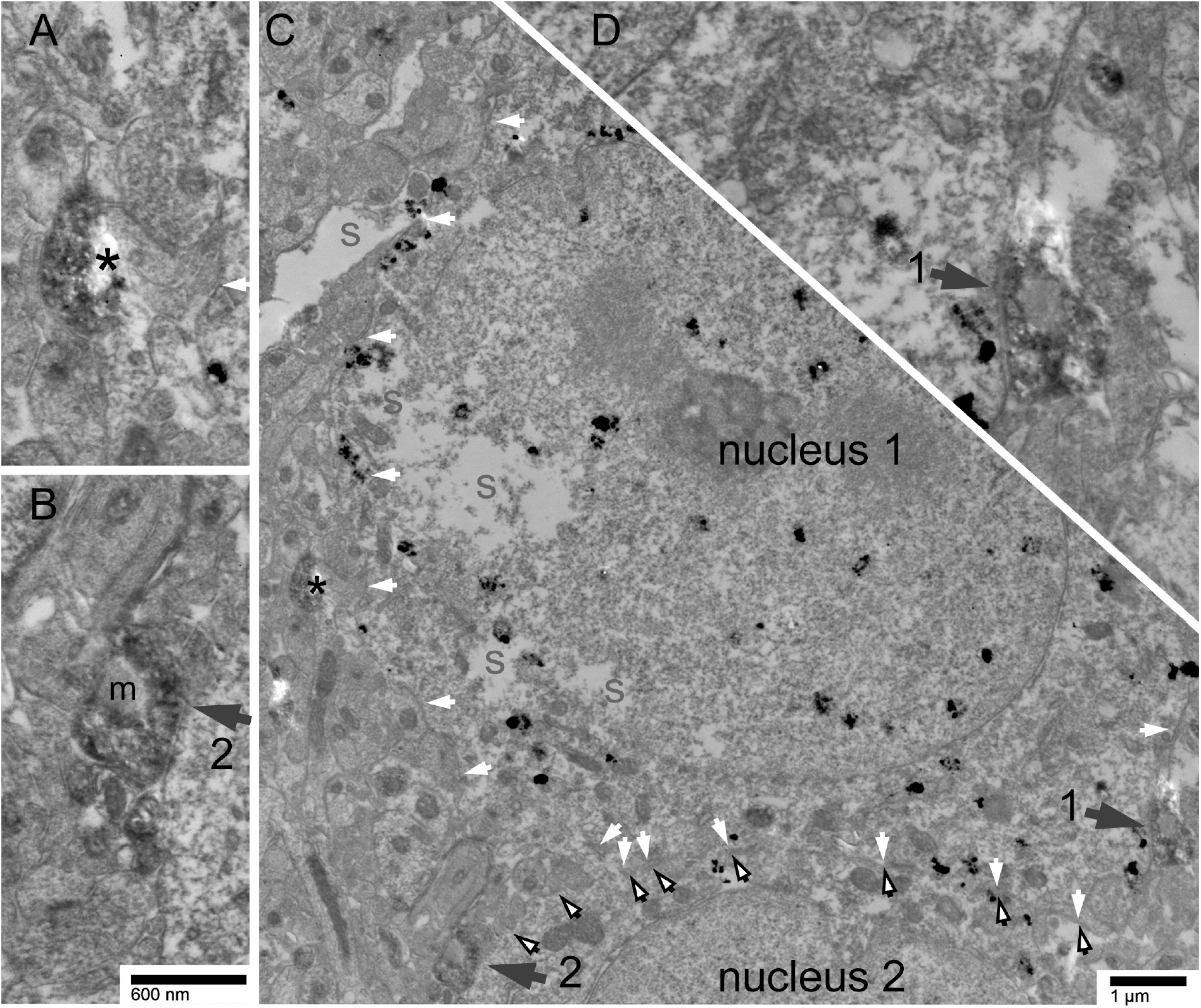
Sparse GABAergic innervation in a pyramidal neuron with mCherry-immunoreactivity from a brain expressing DREADD-Gq among mPFC-DS cells. SIG particles are identifiable by their maximal electron density, approximately 100 nm in size. SIGs are numerous in nucleus 1 and its surrounding cytoplasm, indicating robust mCherry expression (panel B). The plasma membrane of this cell is depicted by white arrows. Another cell body is immediately adjacent to it, indicated by the black-bordered arrows and “nucleus 2” (panel B). SIG particles are evident in the second cell’s cytoplasm as well. GAD-immunoreactivity is evident based on the heavier electron dense precipitate in the cytoplasm that is excluded from vesicle lumens and mitochondria (“m” in the axon terminal forming synapse #2). The two GABAergic innervations evident within this plane of section are indicated by the large black arrows and shown in higher magnification in panels B and D. Arrow #1 and #2 point to a GABAergic innervations of cell 1 and cell 2, respectively. The asterisk (panel A and C) points to a GABAergic axon that is within 600 nm from the plasma membrane of Cell #1 but it interposed by an axo-spinous synapse.

**Figure 8.**
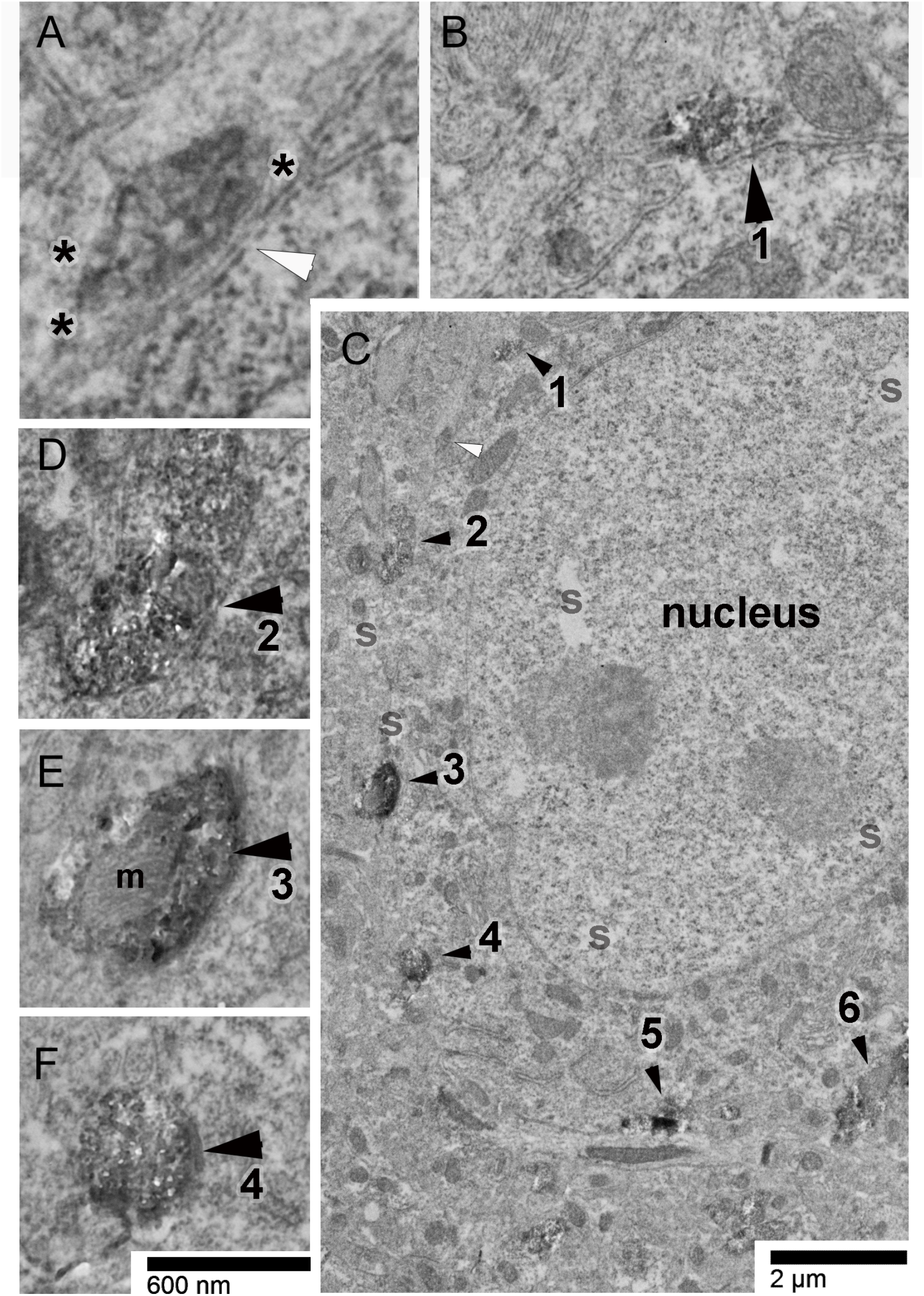
Prevalent GABAergic innervation of a sample pyramidal neuron lacking mCherry-immunoreactivity, from a brain expressing DREADD-Gq among mPFC-DS cells. Numerous GABAergic axon terminals form axo-somatic synapses with this cell body. The image of the cell body was captured near the surface of the vibratome tissue (indicated by the label “S” where no tissue can be seen, panel C) to minimize failure to detect SIG particles reflecting mCherry-immunoreactivity. The lack of SIG particles in this neuron indicates that this neuron is mCherry-negative. The smooth nuclear envelope and absence of GAD-immunoreactivity in the cytoplasm indicates that this is a pyramidal neuron. Six additional GABAergic axo-somatic synapses along the cell body plasma membrane at this plane of section are shown, with details captured at a higher magnification of 30,000x in the surrounding small panels B, D, E and F). GAD-immunoreactivity is evident based on the heavier electron dense precipitate in the cytoplasm that is excluded from vesicle lumens and mitochondria (“m” in the axon terminal forming synapse #2). The open arrowhead at the upper left in panel C and detailed in panel A depicts an example of a GAD+ axon process coursing near but not in direct synaptic contact with the cell body. Instead, this process is enveloped by an astrocytic process, depicted with three asterisks in its cytoplasm. The width of the astrocytic process that is inserted between the axon terminal and the cell body is no more than twice the thickness of the plasma membrane, estimated to be approximately 20 nm. For all small panels, except for panel A, the calibration bar equals 600 nm. For panel A, the same bar equals 300 nm. The 2 μm calibration bar applies to panel C only.

**Figure 9.**
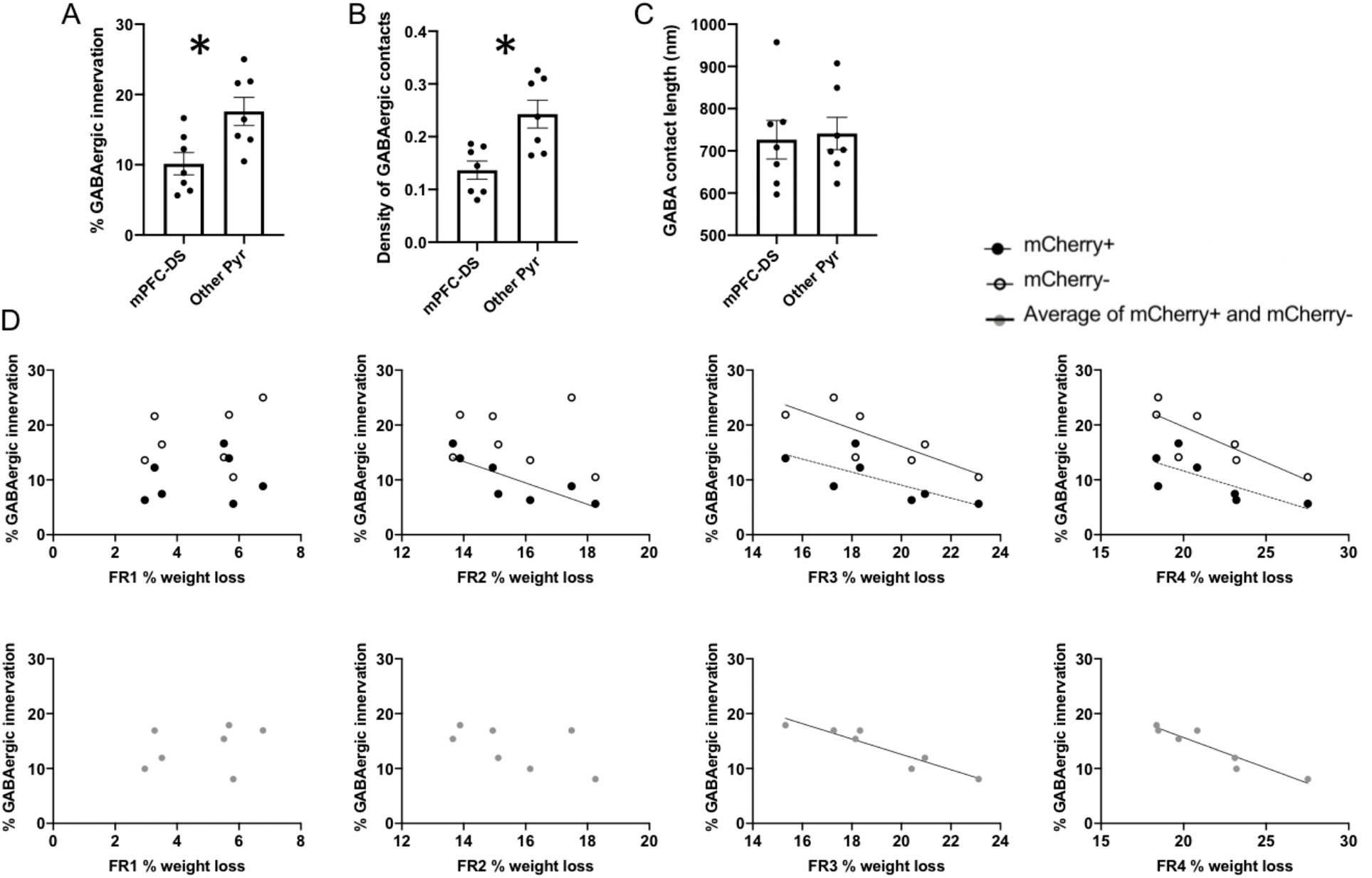
EM analysis of GABAergic innervation of mCherry+ mPFC-DS projecting neurons and unlabeled neighboring cells. **A-C.** Extent of GAD+ terminal synaptic junctions on the somatic plasma membrane of layer 5 mPFC pyramidal cells in the mPFC-DS group were analyzed by EM. mCherry negative cells received less axo-somatic GABAergic innervation than neighboring mCherry negative cells (**A**), with lower density of contacts (**B**) and no difference in terminal size (**C**). **D**. Correlations between percent GABAergic innervation and percent weight loss on each of the 4 days of FR. Significant correlations (p < 0.05) are indicated with a black line of regression, strong trends (p < 0.1) are indicated by dashed lines. Percent weight loss is calculated as the weight at 7pm on the FR day subtracted from the weight at the start of ABA2, all divided by the weight at the start of ABA2.

**Figure 10.**
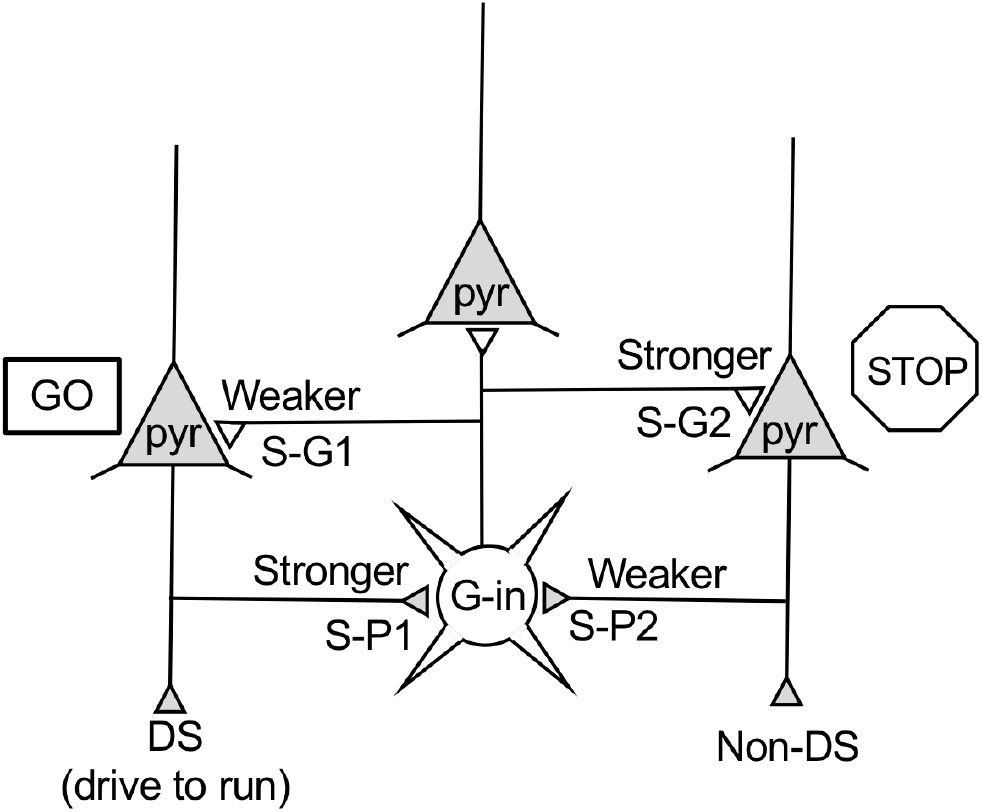
Schematic of mPFC microcircuitry. Pyramidal cells (represented by triangles) that project to DS drive running in the context of food restriction (Fig. 2A). Widening the population of mPFC pyramidal cells to include the non-DS projecting population does not increase running beyond what is observed by driving the DS-projecting pyramidal neurons (Fig. 3A). mPFC-DS cells elicit a stronger response from GABAergic cells (Fig. 5C, synapse S-P1), which in turn innervate non-DS projecting pyramidal cells (synapse S-G2) more prevalently than mPFC-DS projecting pyramidal cells (synapse S-G1, Fig. 9A). This enables the mPFC-DS subgroup to conduct excitatory flow (green light) to elicit food restriction-evoked running, while suppressing the neighboring non-DS projecting pyramidal cells. This model is further supported by correlations between individual differences in FR-evoked running and characteristics of GABA-INs, which were revealed by looking separately at GABA-IN-to-Pyr and Pyr-to-GABA-IN relationships. Axo-somatic inhibition of mPFC pyramidal cells (synapses S-G1 and S-G2) correlated with weight retention in general, but not feeding or running in particular, indicating a protective role for GABA-IN suppression of mPFC pyramidal cells through a combined mechanism of enhanced feeding and suppressed running (Fig. 9D). This effect is in keeping with SalB suppression of running, either by targeting mPFC-DS cells or by targeting the general mPFC pyramidal cell population (Fig. 2C and 3C). Likewise, DREADD activation of the general mPFC pyramidal cell population revealed a negative correlation between evoked activity in GABA-INs and evoked running activity, indicating a protective role of GABA-INs in suppressing FR-evoked running (Fig. 5F). By contrast, when DREADD activation was limited to just the mPFC-DS subpopulation that drives hyperactivity, evoked activity in GABA-INs (synapse S-P1) and evoked running activity correlated positively (Fig. 5F), specifically during FR and not during *ad libitum* food availability. Notably, the stop-light interpretation presented here predicts this positive correlation, as greater mPFC-DS activity produces both greater running (DS output) and greater interneuron activity (synapse S-P1), which preferentially targets non-DS projecting pyramidal cells (synapse S-G2).

### Electron microscopic quantification of GABAergic innervation correlates with individual differences in weight

We also found that axo-somatic GABAergic innervation of both mCherry+ and mCherry-cells correlates negatively with percent weight loss (Fig. 9D), suggesting that GABAergic innervation is protective against the precipitous weight loss that can become fatal in the ABA model. On FR1, mice had the least exposure to FR, and had only lost between 2% and 7% of weight. This measure did not correlate with GABAergic innervation of DS-projecting mCherry+ cells (p = 0.640; R = 0.217), non-DS-projecting mCherry-cells (p = 0.607; R = 0.238), or the average sample of both cell types (p = 0.542; R = 0.281). On FR2, when mice received C21 to drive cell firing in mPFC-DS cells, only the GABAergic innervation upon DS-projecting mCherry+ cells correlated negatively with percent weight loss (p = 0.025; R = −0.815). GABAergic innervation of the mCherry-population did not correlate with weight loss (p = 0.709; R = −0.174). As expected, the average GABAergic innervation of the two cell groups, combined, did not correlate with percent weight loss on FR2, either (p = 0.191; R = −0.560). This suggests that those individuals with strongest GABAergic inhibition of DREADD-driven mPFC-DS cells were the most protected from weight loss following C21 administration on FR2 and that the GABAergic synapses onto non-DS-projecting mPFC cells were relatively less effective in weight preservation. By FR3, percent weight loss trended towards a negative correlation with GABAergic innervation of mCherry+ cells (p = 0.060; R = −0.736), and significantly correlated with GABAergic innervation of mCherry-cells (p = 0.034; R = −0.790), with a significant correlation when GABAergic innervation of the two cell types were combined in the analysis (p = 0.002; R = −0.939). By FR4, which is the day that SalB was delivered, percent weight loss trended towards a correlation with GABAergic innervation of mCherry+ cells (p = 0.069; R = −0.719), and significantly correlated with GABAergic innervation of mCherry-cells, only (p = 0.030; R = −0.802), or with both cell types combined (p = 0.002; R = −0.938). Major findings are summarized in Tables 1 and 2.

**Table 2.**
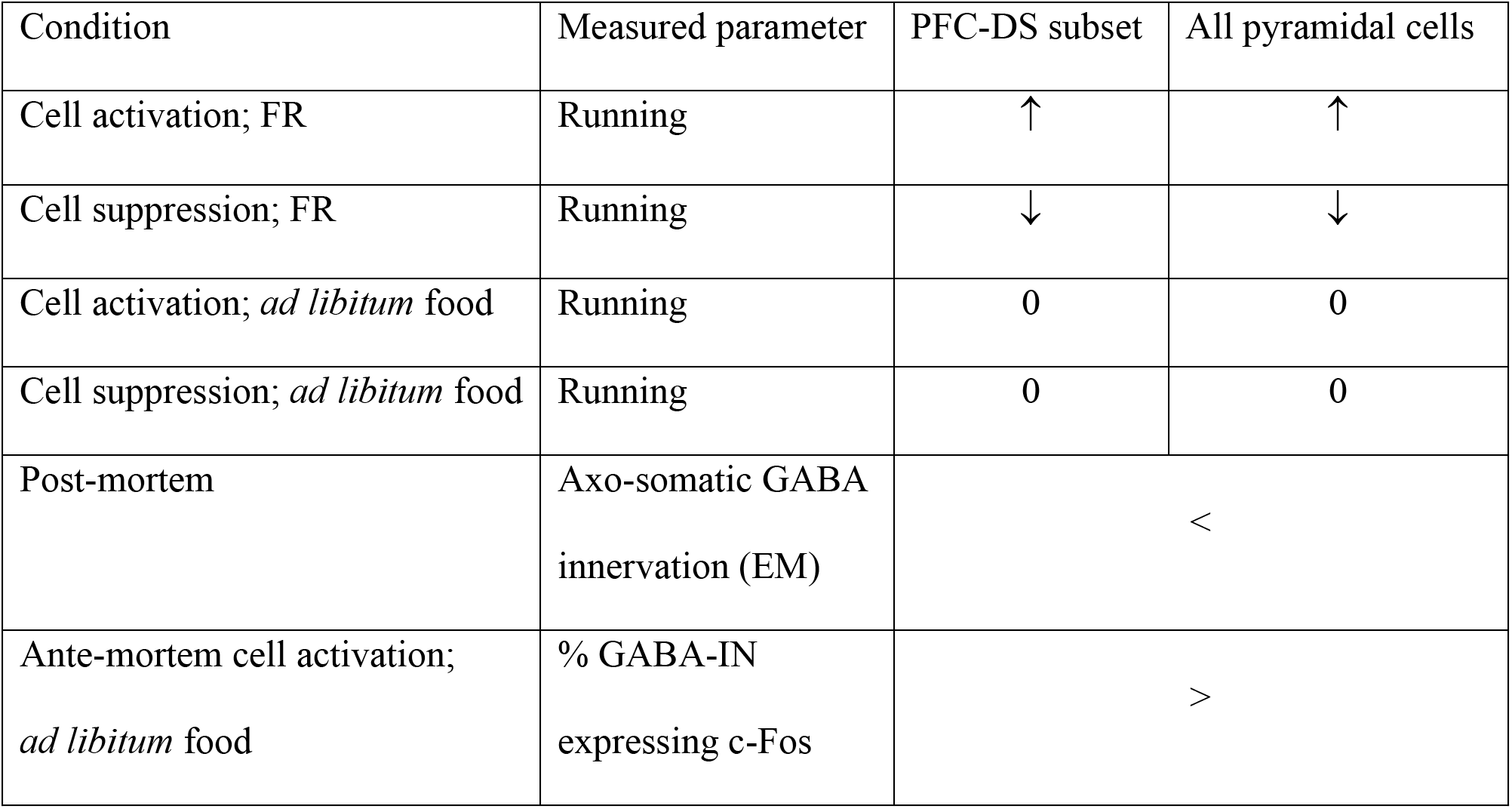
Summary of major results. List of conditions and the observed effects each produced when applied to either the mPFC-DS subpopulation or the general pyramidal cell population. Rows 1-4 reference Fig. 2–3, row 5 references Fig 9; Row 6 references Fig 5C. For rows 5 and 6, < and > signs compare the mPFC-DS population vs the general population of pyramidal cells. Up arrow indicates an increase in the measured parameter, down arrow indicates a decrease in the measured parameter, and 0 indicates no difference.

## DISCUSSION

We demonstrate here for the first time that activation of mPFC pyramidal cells drives an increase in running distance and duration specifically during FAA of FR days, and not during the two hours of food availability, nor post-prandial time bins, and not on days with *ad libitum* food availability in weight-restored mice. This effect was fully recapitulated by driving activity of the mPFC-DS subpopulation of pyramidal cells. Conversely, suppression of mPFC pyramidal cells generally, or targeting mPFC-DS pyramidal cells, reduces wheel running specifically during FR and not on days with *ad libitum* food access. These findings reveal that mPFC activity in general, and the mPFC-DS circuit in particular, do not simply control the drive to run, but rather, control hyperactivity induced specifically by the condition of FR.

While the phenomenon of FAA has been previously linked to activity in the DS (Gallardo et al. 2014), this is the first study to causally implicate the mPFC in FAA. Of all mPFC pyramidal cells, only ~15% project to DS (Gabbott et al 2005), yet we observed a similar increase in running when driving activity in all pyramidal cells, vs the DS projecting subset, suggesting that the mPFC-DS subset drives the observed effects on FR-evoked running. Thus effects could be mediated by any subset of DS cells projecting to specific downstream targets, as well as collateral projections from mPFC to several additional brain regions, including basolateral amygdala, ventral striatum, spinal cord (Gabbott PL et al. 2005).

### Axo-somatic synaptic inhibition preferentially targets non-DS projecting mPFC pyramidal cells

We report here the first evidence that GABA-INs provide less axo-somatic innervation to mPFC-DS projecting cells, vs neighboring mPFC pyramidal cells that do not project to DS. Axo-somatic inhibition is likely to be from fast-spiking parvalbumin-positive cells (PVs), as this subtype is known for strong axo-somatic inhibition of pyramidal cells (Kawaguchi Y and Y Kubota 1997). Fast-spiking PV interneurons preferentially target layer 5 mPFC pyramidal cells with characteristics of PT (pyramidal tract) neurons (thick-tufted apical dendrites, prominent h-current) than of IT (intratelencephalic) neurons (thin-tufted apical dendrites, less prominent h-current) (Lee AT et al. 2014). Striatum is the only subcortical region that receives both PT and IT projections from layer 5 of cortex (Shepherd GM 2013). Based on the axo-somatic GABA input pattern, the mPFC-DS subpopulation could potentially overlap more with the IT than with the PT neurons. IT neurons also project preferentially to D1R-expressing striatal cells (Reiner A et al. 2010), the latter of which are necessary for FAA (Gallardo CM *et al.* 2014). Our data support this notion by revealing that mPFC-DS pyramidal cells also are necessary for FAA.

### Axo-somatic synaptic inhibition of pyramidal cells as revealed by EM is linked to weight loss

Individual differences in axo-somatic GABAergic innervation correlates negatively with FR weight loss, suggesting that increased GABAergic innervation may be protective against weight loss. Interestingly, after C21 delivery on FR2, weight loss correlated negatively with GABAergic innervation only to the DREADD-driven mPFC-DS subset of cells responsible for exacerbated FAA, and not to neighboring pyramidal cells. On FR3 and FR4, weight loss negatively correlated best with GABAergic innervation averaged across all layer 5 pyramidal cells, indiscriminate of their projection pathway. Correlation of GABAergic inhibition with weight loss as a whole, rather than the parameters of running or food consumption individually, suggests that subjects may have taken different combinations of two strategies (increase consumption and/or reduce wheel running) to minimize weight loss.

### Distinction in the microcircuitry of mPFC pyramidal cell subgroups: DS-projecting versus others

Investigation of post-mortem c-Fos activity revealed additional characteristics of the mPFC-DS pathway that distinguish this subset from the general mPFC population. Selectively driving the mPFC-DS subset, vs general mPFC pyramidal cell activity, induced a greater percentage of both mPFC pyramidal cells and GABA-INs to become c-Fos positive. Thus, driving mPFC-DS activity produces two seemingly opposed outcomes: increased FAA running (vs CON subjects) and increased elicitation of GABA-IN activity (vs CaMKII group). When considered in concert with the EM data described above, a new interpretation emerges: the mPFC-DS pathway recruits more GABA-INs, yet receives less axo-somatic feedback inhibition (Figure 10). Thus, mPFC circuitry may produce a stop-light effect, whereby mPFC-DS pyramidal cell firing elicits a strong response from GABA-INs which, in turn, suppresses non-DS projecting pyramidal cells to a greater extent than the mPFC-DS subgroup. This interpretation is consistent with the observation that C21 elicits 5X-greater c-Fos response from the mPFC-DS pyramidal cells, vs the CaMKII pyramidal cells, generally.

### Functional distinctions of the pyramidal cell groups via differential recruitment of GABA-INs

Individual differences in FR-evoked running correlated, not with pyramidal cell c-Fos activity, but instead, with GABAergic response following ante-mortem C21 delivery. For the CaMKII group, C21-driven GABA-IN c-Fos response correlated negatively with running only on FR2, suggesting that individual differences in DREADD-driven GABAergic recruitment accounts for individual differences in running, driven by excitement of those same mPFC pyramidal cells. The negative sign suggests that GABAergic innervation protects against the excessive running that can otherwise become fatal in ABA. Likewise, CaMKII+ pyramidal cell recruitment of GABA-IN, as revealed by c-Fos activity, correlated negatively with weight loss, again suggesting GABA-INs’ protective role.

By contrast, individual differences in GABA-IN c-Fos expression elicited by driving the mPFC-DS subset correlated positively with running, specifically during FR and not during *ad libitum* food availability. Both the positive (for the mPFC-DS) and negative (for the CaMKII group) correlations between GABA-IN c-Fos and running support the stop-light interpretation (Fig. 10), where greater inhibitory control of non-DS cells than of the mPFC-DS cells both support running. By this logic, preferential inhibitory axo-somatic target by PV cells onto non-DS cells allows for preferential execution of mPFC-DS dependent tasks - in our case, FR-evoked running. mPFC-DS pyramidal cell firing is necessary for both reward-related cognitive flexibility (Nakayama H et al. 2018) and goal-directed action selection (Friedman A et al. 2015). Without prior knowledge of the stop-light circuitry revolving GABA-INs, one might expect that mPFC PV cell activation would generally inhibit these mPFC-dependent tasks, but instead, the opposite is reported: PV cell activation improves goal-directed behavior (Kim H *et al.* 2016) and reward-related cognitive flexibility (Sparta DR et al. 2014). By contrast, PV cell activation impairs performance on tasks that have not been shown to require the mPFC-DS pathway, as is the case in working memory, social interaction (Ferguson BR and WJ Gao 2018), and regulation of anxiety (Page CE et al. 2019). Preferential PV innervation of non-DS projecting pyramidal cells may explain why PV activation impairs tasks dependent on non-DS projecting cells, while enhancing mPFC-DS dependent tasks. Notably, the majority of works cited here (Sparta DR *et al.* 2014; Kim H *et al.* 2016; Ferguson BR and WJ Gao 2018; Nakayama H *et al.* 2018) use FR to reduce body weight to ~80-90% of baseline to elicit behavior, which is similar to the percent reduction we observed on FR2. Our findings indicate that FR used in these reward-related cognitive flexibility and goal-directed action selection tasks could have enhanced the engagement of mPFC-DS pathways through increased activation of mPFC PV neurons.

### mPFC-DS pathway in decision-making

In the context of severe FR, an organism is presented with the decision to either 1) reduce caloric output (low cost) yet remain in the low-resource region or 2) increase caloric output (high cost) and run to a region that may provide increased resources (potentially high reward). Friedman et al found that the mPFC-mDS striosome pathway is necessary for cost-benefit analysis under an approach-avoidance conflict (Friedman A *et al.* 2015) and that chronic stress increased suboptimal high cost/high reward choices via increased excitation of mPFC-driven DS cells (Friedman A et al. 2017). Our ABA mice experienced both the chronic stress of single-housing as well as the stress of FR. This environmental stress may likewise have opened the gate for C21 modulation of mPFC-driven DS pathway, tipping the balance towards the high cost/high risk decision to run under conditions of low caloric availability. Conversely, closure of the DS gate during the days of *ad libitum* food, free of chronic stress, may explain why C21 could not tip the balance towards the animals’ decision to run on days of recovery.

### Clinical relevance

The effect of extreme FR in humans is most extensively studied in the context of Anorexia Nervosa (AN), a disorder of self-imposed FR. Nearly all subjects with AN exhibit compulsive hyperactivity, which contributes to the severity of the disorder (Sternheim L et al. 2015). In AN, DS is more engaged in making food choices, as compared to healthy controls, and connectivity between mPFC and DS is stronger when subjects choose maladaptive low calorie vs high calorie foods (Foerde K et al. 2015), indicating a possible role for the mPFC-DS pathway in AN. Cognitive rigidity and tendency towards stereotyped behaviors, including compulsive running, as well as heightened anxiety are hallmarks of AN (Walsh BT 2013). GABA_A_ receptor agonists, used to treat anxiety, are not effective for treating AN, which currently has no FDA approved pharmacological treatment (Frank GK and ME Shott 2016). Our data suggest this could be because GABAergic innervation targets primarily non-DS projecting cells, rather than the mPFC-DS pathway that mediates FR-evoked hyperactivity.

## ACKNOWLDGEMENTS

Acknowledgement of assistance: We gratefully acknowledge Ishan Handa, Sabrina George, and Rose Temizer for their role in data collection. We gratefully acknowledge the NYU Office of Veterinary Resource staff and Claudia Farb for their technical support and advice.

